# Beyond Ramachandran distributions: quantifying secondary structure effects on amino acids and their pairs in the full torsional space

**DOI:** 10.64898/2026.10.09.758016

**Authors:** G. Carrière, C. H. Robert, F. Cazals

## Abstract

The Ramachandran diagram has long served for understanding protein secondary structure elements (SSEs), despite the multitude of factors contributing to their occurrence. Starting from recently introduced circula-based multivariate torsion-angle distribution (CBMD) mixture models – accommodating covariance structure in the full torsion-angle space of amino acids, we define continuous densities for the 20 amino acids and 400 amino-acid pairs in dimension up to 14, conditioned by SSE labels. Projecting these higher-dimensional distributions onto the classical Ramachandran diagram reveals how secondary structures in the protein universe emerge from amino-acid-dependent conformational preferences reflecting influences beyond backbone dihedrals alone. Our analysis links the traditional geometric description of local protein structure to modern statistical models of the full conformational space of amino acids and amino-acid pairs, while providing a framework for generating protein conformations consistent with any given secondary structure environment – a feature that remains difficult to enforce explicitly in current AI-based generative models.

## 1. Introduction

### 1.1 Previous work

In courses presenting the topological structure of the polypeptide chain, the Ramachandran diagram, introduced by G.N. Ramachandran, C. Ramakrishnan and V. Sasisekharan in their seminal 1963 paper [1], typically figures prominently alongside the *α*-helix and *β*-sheets — the original secondary structure elements discovered a decade earlier by Pauling and collaborators [2, 3]. These fundamental biophysical concepts allow bridging the gap between the realm of physics and potential energy models on the one hand, and higher-level abstractions on the other.

The connexion to energy models may be illustrated by quotes from textbooks, such as “*steric clashes restrict the number of conformations available to the polypeptide chains*” [4](Chapter 1), or “*Not all values of φ and ψ are accessible to the peptide backbone*… … *certain values of φ and ψ bring atoms of the backbone into such close proximity that the repulsive component of the van der Waals energy prevents the atoms from coming any closer*” [5](Chapter 4). The link to higher-level abstractions, for its part, is seen in the widespread superimposition of secondary structure elements (SSE) labels onto scatter plots of the dihedral angles *φ* and *ϕ* which embody the Ramachandran diagram.

The joint presentation of Ramachandran diagrams and SSE has both a merit and an intrinsic difficulty. The clear merit is to provide a qualitative and intuitive bridge between these two closely related notions. The difficulty is that energy models and SSE labels are specified in related but different ways. On the one hand, values of (*φ, ψ*) angles can be used to infer distances between backbone and side chains (*C*_*β*_) atoms of two consecutive amino acids, from which potential energy consequences may be computed [6, 7]. On the other hand, SSE are structural patterns requiring a local environment beyond the isolated amino acid backbone, and are discrete in nature. This discreteness inherently imposes some arbitrariness in defining boundaries of the regions associated with SSE labels in Ramachandran space. Formal methods such as DSSP [8, 9] use criteria on hydrogen bond topology to assign SSE labels (8 possible labels) to amino acids. Other alternatives may be considered [10, 11], notably `STRIDE` [12] (7 possible labels), which combines hydrogen bond topology and Ramachandran angles.

This complexity has motivated continuous work on these topics. Delicate analyses of atomic repulsions and attractions have been elaborated to explain the geometry of excluded and occupied regions [6, 7]. These analysis used a combination of methods, including model peptides, molecular dynamics, and PDB data analysis. Significant efforts have been undertaken to refine the definitions of Ramachandran regions [13] using complementary criteria. While it has long been acknowledged that Ramachandran regions cannot uniquely determine secondary structure — both notions are defined in different spaces in the first place — superimposing SSE labels onto Ramachandran diagrams remains common practice. Indeed, the challenge of assessing structure reliability while accompanying the relentless increase in high-resolution protein structures deposited into the PDB has spurred refinement of criteria for labeling SSE [9] and development of new tools based on Ramachandran+SSE plots [14]. This ongoing effort may be represented by superimposing the boundaries of excluded/admissible regions of the Ramachandran plot along with the labels of regions corresponding to SSE (Fig. 1(Top)), and comparing these to sterically-excluded regions (Fig. 1(Middle)).

**Figure 1.**
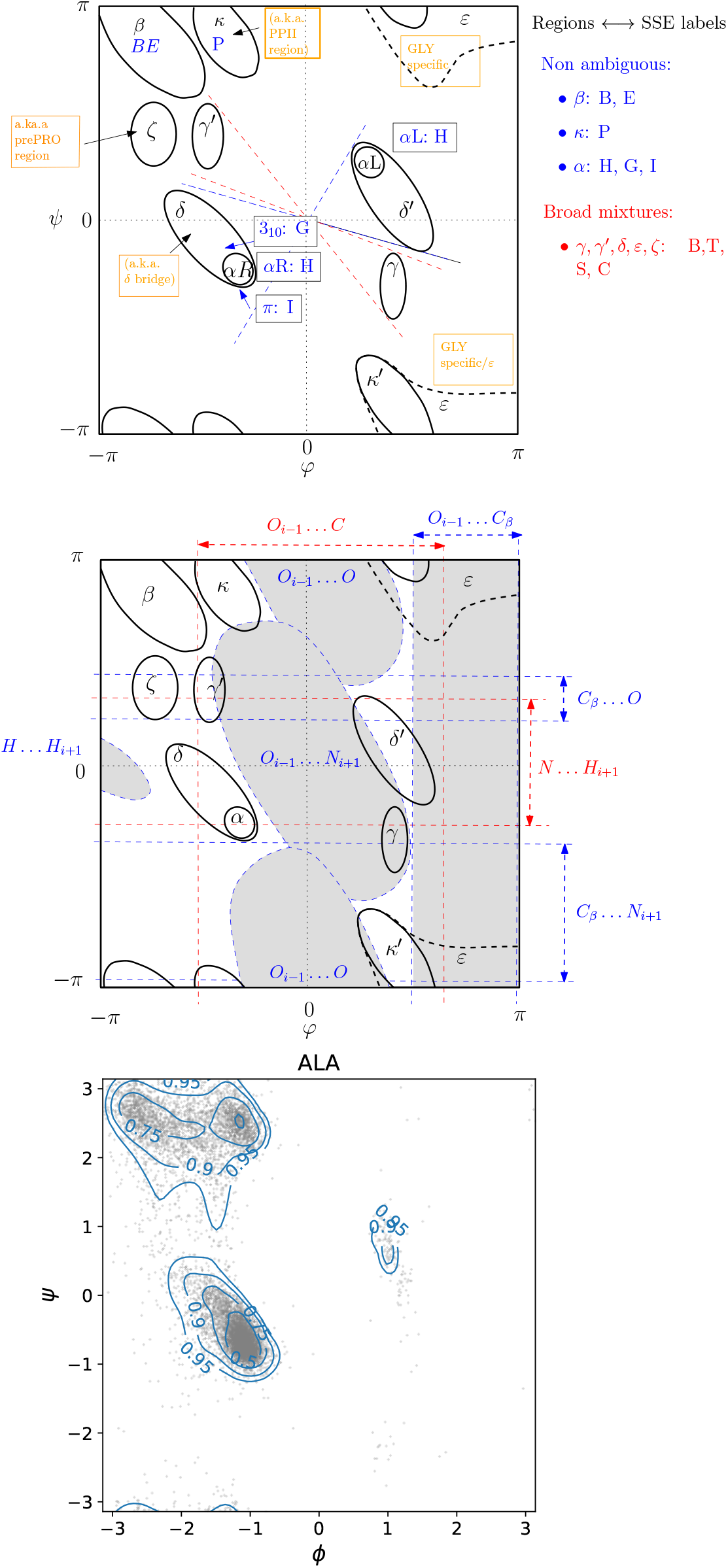
Ramachandran diagrams: idealized regions versus probability density (here ALA). **(Top)** Region nomenclature of A. Karplus and coworkers [13]. Dashed lines indicate symmetries with respect to (0, 0). **(Middle)** Sterically-excluded regions (in gray) from Brasseur et al. [6] (GLY, PRO, pre-PRO excluded). For GLY and pro-PRO, see [7]. **(Bottom)** Generated samples and credible region contours from an amino acid agnostic density model by Carrière et al. [18] estimated on a corpus of ALA observations, irrespective of SSE. See details in Sec. S1.

Stepping away from Ramachandran strictly speaking, other important connections have also been made. One is the definition of rotamer libraries providing (discrete) rotamer frequencies given a (continuous) Ramachandran position [15]. More recently, the *cross-landscape* diagram reveals localized regions in the diagram (*ψ*_*i*_, *φ*_*i*+1_) reflecting conformational and dynamic properties inherent to the rigidity of the peptide bond [16].

### 1.2 Contributions

We establish quantitative bridges between Ramachandran regions defined in [13] and SSE labels by presenting probability densities for backbone and sidechain torsion angles in individual amino acids and in amino acid pairs, conditioned on given secondary structures. For a single amino acid, these densities are defined over the flat torus T^*d*^, namely from T^2^ (GLY) to T^7^ (ARG). For pairs of a.a., we condition our densities in T^4^ to T^14^ by pairs of SSE labels.

We use these conditioned densities for two purposes. On the generative side, our densities are plain generative models of amino acid torsion angles given secondary structures. On the analytical side, these densities enable formal decompositions of Ramachandran space based on credible regions capturing a specified fraction of the density, which is instrumental to assessing the coherence with the Ramachandran regions. Further, they yield a direct quantitative analysis of the relationship between rotameric distributions and SSE labels.

## 2 Methods

### 2.1 Overall approach

Let ℒ = {1, …, *S}* represent a dictionary of SSE labels, used to assign a label to every amino acid in a structure.

Our objective is to derive SSE specific mixture models of amino acids, that is *p*(*x*|*j*) with *x ∈* T^*d*^ and *j ∈ ℒ*. A mixture model is constructed with *K* components, each defined by a positive weight *w*_*k*_ (with ∑*w*_*k*_ = 1) and a density *f*_*k*_, so that

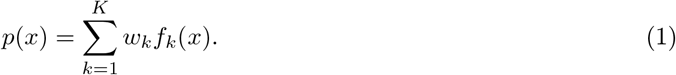

A straightforward strategy consists of partitioning a dataset 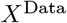 of amino acid observations into SSE-specific subsets 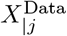 and independently estimate a density for each subset. However, this requires estimating *S* different densities for a given SSE labeling scheme. Furthermore, specific SSE subsets may be represented by few observations, making independent estimation prone to overfitting.

Instead, we build upon SSE-agnostic mixture models from the literature, which provide a complete representation of the amino acids conformational space. With *M* an SSE-agnostic mixture model of *K* components, assume that the conformational patterns of individual SSE are represented among the components of *M* . We specialize M by measuring the degree of association of each of the *K* components to each label *j* to obtain a new reweighted mixture model. Consequently, the component densities *f*_*k*_(*x*) and number of components *K* remain unchanged, while only the mixture weights *w*_*k*_ are modified to obtain the SSE-specific densities *p*(*x*|*l*).

This strategy offers two advantages:

1. With *c*_*f*_ the cost of evaluating a mixture model component *f*_*k*_(*x*), the complexity of conditioning a mixture model on a subset 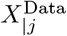 of size *N* is *O*(*NKc*_*f*_). For comparison, each iteration of the classical EM algorithm for estimating a mixture model with *K* components contains an E-step with this same complexity. Conditioning is therefore orders of magnitude faster than a from-scratch estimation of an SSE specific mixture model, which additionally involves estimating the correct number of components [17, 18].
2. The SSE conditioned mixture models rely on more robust component estimates than models estimated independently on small SSE subsets.

### 2.2 Baseline models and datasets

#### Mixture models for torsion angles

We study distributions on the flat torus T^*d*^ coding protein torsion angles, which fall in the realm of Directional Statistics [19, 20].

Two sets of such models have been proposed in the literature. First, the PhiSiCal approach [17] defines 20 mixture models, one per amino acid type, in spaces from T^2^ to T^7^. These models use components defined as products of univariate von Mises distributions — analogues of Gaussian distributions for periodic spaces, and contain no covariance structure. Second, the related circula-based multivariate distribution (CBMD) mixture models of [18], whose component form generalizes that of PhiSiCal to accommodate a covariance structure on torsion angles. This approach provides 20 + 400 + 1 = 421 mixture models: 20 models for individual amino acids, from T^2^ (GLY) to T^7^ (ARG); 20 *×* 20 models for all possible amino acid pairs, from T^4^ to T^14^; and one additional model estimated from backbone consecutive (*ϕ, ψ*) pairs regardless of the amino acid types (T^4^).

Both types of models are estimated from the PDB50 corpus by minimizing Message Length [21], to strike a balance between expressivity and model complexity. In the following, we use the CBMD models as our SSE-agnostic models, due to their superior performance and the possibility of studying amino acid pairs. An example of 10,000 generated points for ALA using the CBMD model (irrespective of secondary structure) is shown in Fig. 1.

#### Protein dataset

The aforementioned CBMD models have been estimated using the PDB50 corpus of proteins [17], a set of 38895 structures with non-redundant amino acid sequences from the Protein Data Bank [22], containing only structures with an R-factor cut-off at 0.3 and resolution cut-off at 3.5 Å or better. To ensure the SSE subsets used for specialization are well represented by the CBMD models, we build them directly from the same PDB50 corpus.

#### SSE labels

We use the `DSSP4` [9] algorithm to assign each amino acid observation in PDB50 one of 9 different labels. `DSSP4` is primarily based on hydrogen-bond patterns rather than backbone torsion angles, and explicitly identifies polyproline helices, a critical topic for analyzing intrinsically disordered regions. We note that `STRIDE` additionally exploits Ramachandran propensities during assignment [12], making it less suitable for studying the statistical relationship between backbone torsion angles and secondary-structure labels.

`DSSP4` provides eight labels, to which we append a ninth one denoted C. These labels relate to Helical structures G = 3_10_-helix, H = *α*-helix, I = *π*-helix, and P = *κ*-helix (poly-proline II helix); Beta structures B = residue in isolated *β*-bridge and E = extended strand or participating in a *β* ladder; and Other forms S = bend, T = hydrogen-bonded turn, and C = remaining cases/coil.

To specify a CBMD model for an amino acid to a particular secondary structure, we provide one of these 9 labels. To specify a model for a pair of a.a.’s, we provide a pair of labels taken from the Cartesian product of the nine labels.

### 2.3 Conditioned mixtures as reweighted mixtures

In the following, we state the analytical expression of the mixtures fitted on torii ranging from T^2^ (GLY) to T^14^ (ARG-ARG). The reader is referred to Section S2 for the proofs.

#### Conditioned densities

We fit probability densities as reweighted mixture models with components *f*_*k*_. We define the vector (*m*_*k*1_, …, *m*_*kS*_)^T^ of responsibilities of the labels for the k-th component. Note that ∑_*j∈ℒ*_ *m*_*kj*_ = 1, *∀k*. Using these, one has:

##### Proposition. 1

*The conditional density with respect to label j satisfies:*

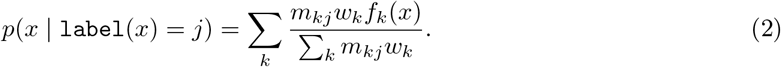

#### Densities and credible regions

We illustrate densities using credible-region contours. For a density *f* and a mass fraction *t ∈* (0, 1), define the cutoff *c*_*t*_ so that 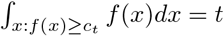. The *credible region* or *highest density region* at threshold *t* is the contour plot/level set surface associated to *c*_*t*_.

#### SSE label uncertainty in Ramachandran space

We investigate the link between Ramachandran coordinates and secondary structure by estimating the entropy of SSE labels depending on the Ramachandran position. To this end, we bin the Ramachandran space in a grid and estimate the SSE label entropy in each grid cell – Eq. 14.

#### Jensen-Shannon divergence between conditional distributions on SSE

We wish to assess the dependency of torsion angles in a given amino acid to the SSE label, either to all torsion angles or to side chain angles only. We do so by estimating the Jensen-Shannon divergence *D*_JS_ between our SSE-conditioned models. That is, with *j*_1_ and *j*_2_ two labels, we compute *D*_JS_ (*p*(*x* | *j*_1_)*∥p*(*x* | *j*_2_)) – Eq. 15 and Eq. 17. With logarithms in base *e*, 0 *≤ D*_JS_ *≤* ln 2 [23].

## 3 Results

All conditioned model parameters are publicly available on the Structural Bioinformatics Library website^1^, along with python code to load and use the models.

### 3.1 Ramachandran regions versus SSE labels

#### Comparison via credible regions

For each amino acid, we computed the nine SSE-conditioned probability densities (one for each `DSSP4` SSE label) from eq. 2. These probability distributions include all relevant heavy-atom torsion angles in the amino acids of the PDB50 dataset, and constitute smooth functions describing the resulting point clouds on the torus T^*d*^ up to dimension *d* = 7. Using this model, we can calculate credible regions enclosing any desired probability mass fraction of the joint distribution of the torsion angles of any given amino acid.

We obtain one plot per amino acid (Fig. 2, selected cases; Fig.S1, all amino acids), within which each SSE-conditioned density is represented by its credible region at *t* = 0.75; *i*.*e*. the contour that captures 75% of the total density. For the sake of comparison, each plot also depicts the reference Ramachandran regions (from Fig. 1(Top) and [13]). As a quality check, we also co-locate contours with data points in Fig. S2, and provide label-specific plots with credible contours at various levels (G to C, Fig. S8 to Fig. S16).

**Figure 2.**
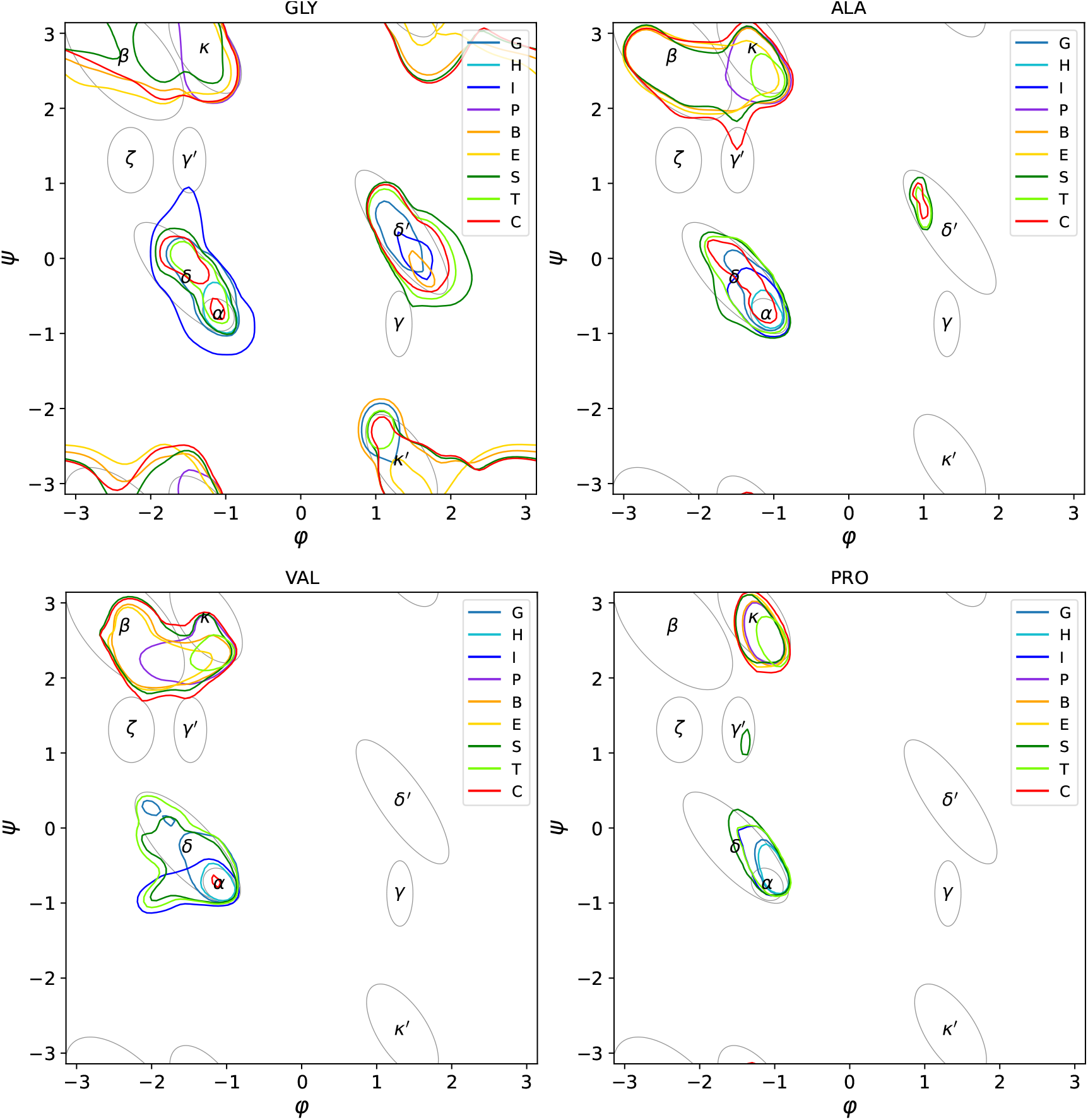
Secondary-structure-conditioned densities superimposed on Ramachandran regions highlight their dependence on amino-acid identity and secondary structure. (Selected examples; see all densities in SI.) Credible region contours of conditioned densities at 75% level.

The resulting plots yield, per amino acid, a statistically substantiated definition of the SSE-defined regions (Fig. 2 versus Fig. 1). The correspondence between reference Ramachandran regions and SSE-conditioned probability densities clearly varies substantially depending on the amino acid type.

The credible regions illustrate the complexity inherent to SSE labels (Fig. 2 and Fig.S1):

- The label H/*α*-helix has a well-defined region. But those of label G/3_10_-helix and label I/*π*-helix are more fuzzy, with (in general) a significant overlap with labels S, T and C.
- The label I/*π*-helix has a specific pattern for a.a. ILE/VAL/THR, with an outgrowth towards negative values of *φ*.
- The labels B and E coding strands are highly coherent.
- Labels S/T/C are in general mingled in three broad regions, namely *α* + *δ, β* + *P*_*II*_ and *δ*^*′*^ . This observation hints at difficulty of assigning SSE labels in these regions.
- The label C displays a specific pattern for ASN, ASP, CYS, HIS, PHE, TRP, TYR, as the curve hits the regions *ζ* and/or *γ*^*′*^ .

Of particular interest are the discrepancies between reference Ramachandran regions and credible contours:

- *P*_*II*_ region: for ARG/LYS/MET in particular, the densities escape their traditional region, yielding a significant overlap with the region *β*.
- region *ζ*: HIS/PHE have more mass around *ζ* region than the other amino acid types
- region *γ*^*′*^ : ASP/LEU have more mass than the otheramino acid types

With a focus on individual amino acids:

- PRO has a well resolved diagram, with all credible curves concentrated in the regions *α* + *δ* and *P*_*II*_.
- ASP has a distinct lack of “C” (coil) observations in the top left of the beta regions not present in other amino acids

Finally, credible curves also shed light on possible discrepancies between Ramachandran regions and physical constraints, in particular the fact that the region *γ* is located inside/on the border of the exclusion region associated with the pair *O*_*i™*1_ … *N*_*i*+1_ (Fig. 1(Middle)). The region *γ*, centered around (*φ* = 65^*°*^, *ψ* = *™* 70^*°*^), is relatively rare and corresponds to a backbone H bond between *CO*_*i*_ and *NH*_*i*+2_ [13, 24]. This H bond gets the `DSSP4` label T, which covers more broadly H bonds *CO*_*i*_ and *NH*_*i*+*n*_ not already assigned to a helix.

We first note that the regions defining the steric map are *best -fit* curves which do contain samples in the *γ* region [6](Fig 5A). These steric regions were defined by a combo of methods including a model peptide, molecular dynamics, and PDB based analysis. By inspecting credible contours for all `DSSP4` labels (G to C, Fig. S8 to Fig. S16), we note that the *γ* region is seldom populated, with contributions from the pairs (B; GLY), (S; GLY, HIS, ILE, TRP, VAL); (T; GLY). Moreover, the case of GLY is peculiar, since the absence of *C*_*β*_ relaxes the entire structure [6]. Altogether, the nature of the steric curves fit using a model peptide, the rarity of these conformations, their presence in an entire protein rather than a model peptide, and the prevalence of glycine resolve the apparent contradiction.

#### Comparison via the entropy of labels

The previous examples illustrate difficulties in reconciling Ramachandran regions and SSE labels. One such complication arises when more than one type of SSE may be associated with a given point in the *ϕ*_*i*_, *ψ*_*i*_ plane. This effect can be quantified in terms of maps of the SSE label uncertainty, or entropy, throughout the plane for each amino acid (Fig. S3). In these plots, the darkest blue color indicates regions with high certainty, whereas turquoise and green-yellow reflect increasing ambiguity for SSE contributions. As expected, the least ambiguous Ramachandran regions are the *α*-helix and much of the *β* regions, with a clear exception for PRO, and for ASN/ASP which show more *β*-region uncertainty. At the other end of the scale, the uncertainty tends to be concentrated in the *κ*-helix region and in the *South-West* of the *δ* bridge region. One can also compare the entropy of individual labels on the Ramachandran map (Eq. 14 and Fig. S3), and their credible regions (Fig. S8 to Fig. S16). Inspection of the credible contours in individual maps shows a clear contribution of SSE labels {S, T, C} in addition to {G, H, I} (resp. {B, E, P}) near the *δ* (resp. *κ*) region. Visually, this observation corresponds to an entropy value near ln 6 *≈* 1.79 – comparable to the entropy associated with a uniform distribution over six labels.

### 3.2 Backbone torsion angles as prime determinants of SSE labels

On a per amino acid basis, we study the influence of SSE labels on amino-acid torsion angle distributions by computing the Jensen-Shannon divergence between any two labelled distributions. We performed this comparison for labelled distributions in the full torsion angle space *i*.*e*. for backbone and side-chain angles (Fig. S4). Repeating the analysis using only the backbone angle subspace results in very similar divergences values. Indeed, considering all SSE/amino acid types (20 *×* 9(9 *™* 1)*/*2 = 720 values), we obtain the following values for the difference (min=-0.048, mean=-0.0005, median=0.00, max=0.063). This indicates that the JS divergences are driven principally by the backbone torsion angles. We then performed the JS divergence comparisons using only the side chain angles subspace (Fig. S5), to investigate the impact of SSE change in side chain torsion angle distributions independently from the backbone.

Focusing on the full conformational space (Fig. S4), the largest divergences appear between the {G, H, I} and the {P, B, E} models. This should be expected, as they include the *α* and *β* regions, which are classically distinct and make the largest contributions to secondary-structure in proteins. The label T (turns) bears similarities to those of helices *i*.*e*. {G, H, I}, and notably differs from the labels {P, B, E}. The label S (bends) distributions resemble most those of the extended (E) class. Finally, label C is little informative but shows mild similarity to the labels {P, B, E, S}.

Focusing on the side-chain torsion angle space only (Fig. S5), the differences in JH divergences between SSE-conditioned distributions are smaller (less than 0.25), so the divergences in this Figure are presented relative to that value using a different colormap to reflect this change. For all amino acids, one sees that the H label is the one that influences side chains the most, followed by the G label. This effect is more or less visible, for example higher in {PHE, TRP, TYR}, lower in {CYS, LYS, SER}. These observations do not depend upon the number of *χ* angles. ASN and ASP share a pattern where side chains angles are influenced by whether the SSE label is B (*β*-bridge) or C (unknown). For ILE, the G label results in side chain angles that uniquely differ from other labels, including other alpha labels {H, I}. This pattern is also weakly displayed by other alpha labels {H, I, S, T}. Interestingly this does not appear on LEU, but appears in VAL (although less importantly).

Overall, these analyses provide a quantitative assessment of the relationship between torsion angle distributions and secondary structure element (SSE) labels. The results identify backbone torsion angles as the primary determinants of SSE assignment, while side-chain angles play a secondary role, exhibiting amino acid and SSE-specific patterns.

### 3.3 Amino-acid pair conformations conditioned on SSE labels

We next consider conditioned densities in the cross-landscape associated with two consecutive amino acids (*amino dominos*) [16]. The cross landscape is typically presented as a row of three plots (*φ*_*i*_, *ψ*_*i*_), (*ψ*_*i*_, *φ*_*i*+1_) and (*φ*_*i*+1_, *ψ*_*i*+1_) — that is, the Ramachandran plots for the two amino acids at *i* and *i* + 1 with the cross plot (*ψ*_*i*_, *φ*_*i*+1_) in between. Omitting the SSE label C, one can thus define a total of 20 *×* 20 *×* 8 *×* 8 = 25, 600 amino-acid pair probability densities conditioned on SSE classes.

#### Asymmetry in SSE transitions

We employed these amino-acid pair densities to explore transitions from one SSE to another. For this analysis, we reduced the combinatorial complexity by (i) considering all a.a. types together, and (ii) grouping the SSE labels into four classes, namely helices {*G, H, I}*, strands {*B, E}*, the *κ*-helix {*P}*, and bend-turns {*S, T}*. This yields 16 possible permutations of SSE structures of amino-acid pairs. We then present their cross-landscape densities in two Figures:

- Fig.S6, {*-Helix, *-Strand, *-*κ*Helix, and *-BendTurn}, describes the entry from an a.a. of any SSE class (represented by *) into an a.a. of a single specified SSE class, with that class arranged from top to bottom in the figure.
- Fig.S7, { Helix-*, Strand-*, *κ*Helix-*, and BendTurn-* }, refers to the exit from the same specified SSE class into any SSE class.

For a case study, we focus on the entry-exit of the Helix SSE class (Fig. 3). In each of these cross landscapes, the different SSE classes of the a.a. indicated by * in the a.a. pair are represented by their 75% credibility regions to facilitate comparison. Also note that we have included self-transitions (from a given SSE to the same SSE) for completeness.

**Figure 3.**
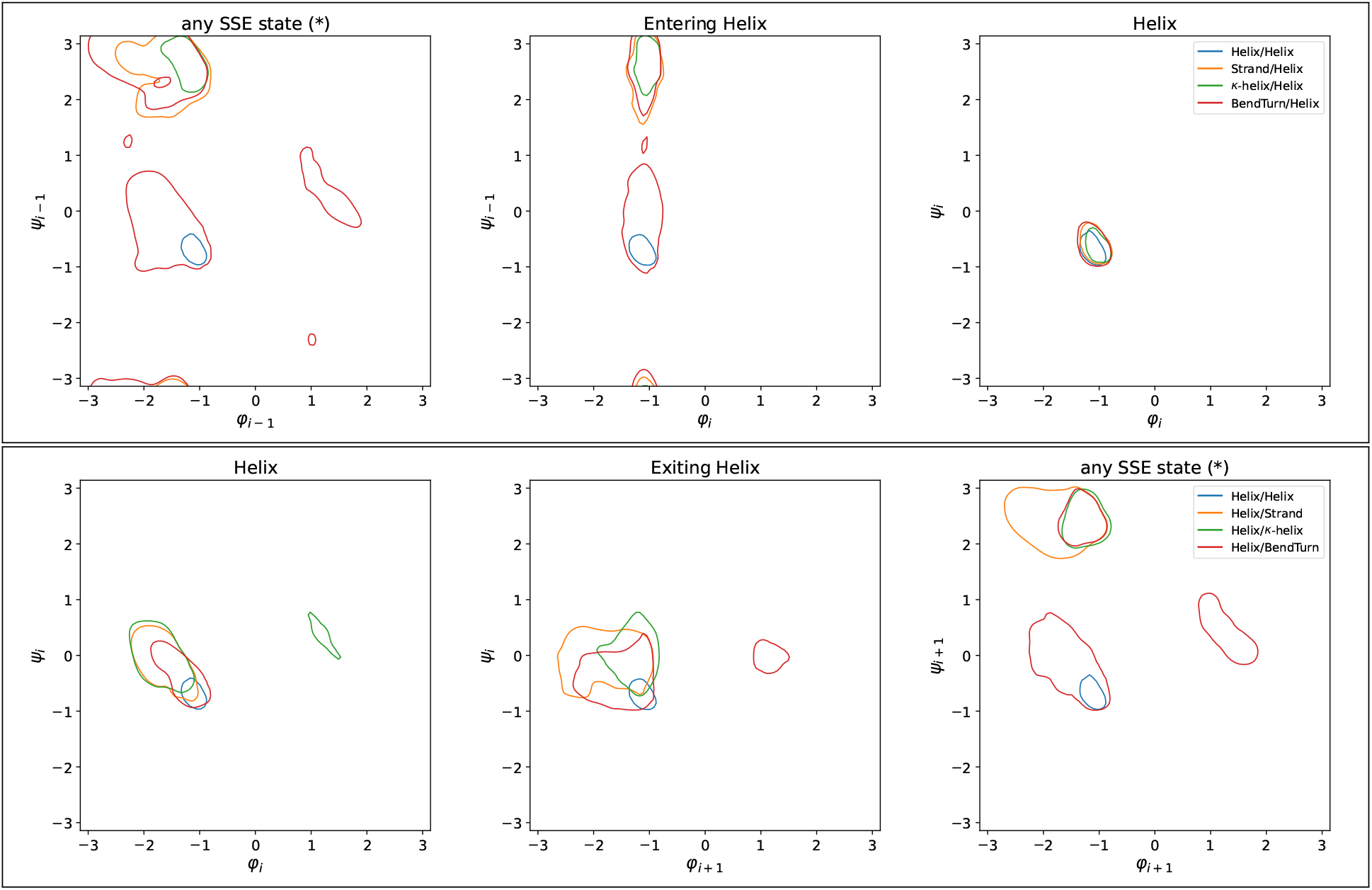
Entering/Exiting a helix: densities of specialized submodels in amino acid pairs illustrate the asymmetry of SSE entries and exits. All amino acids considered. Helix combines the {‘H’,’G’,’I’} DSSP labels, Strand combines {‘B’,’E’}, *κ*Helix is {‘P’}, and BendTurn combines {‘S’,’T’}. Credible region contour at 75% level.

We note that there is considerable asymmetry in these landscapes, in that credible regions Label1-Label2 differ from those tagged Label2-Label1. Generally, the entered (*i*.*e*., second) residue is less affected by the SSE of the exited (first) residue than vice versa (compare Fig.S6 (Right column) to Fig.S7 (Left column)). As an illustration, let us compare the two transitions *-Helix versus Helix-*. In the *-Helix case, the entered residue (helix) is only moderately affected by the SSE of the exited residue, as its corresponding Ramachandran projection remains focused in the *α* region in all cases (Fig. 3(First row, Third column)). In contrast, in the Helix-* case, the helical first amino acid’s probability density is less concentrated in the *α* region. Indeed, the credible regions for this residue in SSE environments Helix-Strand and Helix-*κ*Helix, while assigned as helical, are seen to extend into the *δ* region more typically hosting B/E/P labels (Fig. 3(Second row, First column)).

Transitions between SSEs in consecutive amino-acids have previously been studied using angles (*ϕ*_*i*+1_, *ψ*_*i*_) [25] or by clustering backbone stretches involving six atoms – the amino-domino approach from Rosenberg et al. [16]. As an illustration of the differences from our approach, compare Fig. 3(Top row), calculated using our SSE-conditioned densities, to clusters 9 and 10 in Fig. 4 of reference [16] (NB: in that figure, the hashing patterns for amino acids referred to as *k* and *k* + 1 appear to have been erroneously swapped.) The conditioned densities provide quantitative versions of the clustered observations, for all types of transitions and all amino acids. They effectively replace the scatter plots in Fig. 4 of [16] for a given cluster by a conditioned probability density in dimension up to *d* = 14. Further, they can be used quantitatively in detailed structural analyses, as shown for a particular case of helix capping mechanisms.

**Figure 4.**
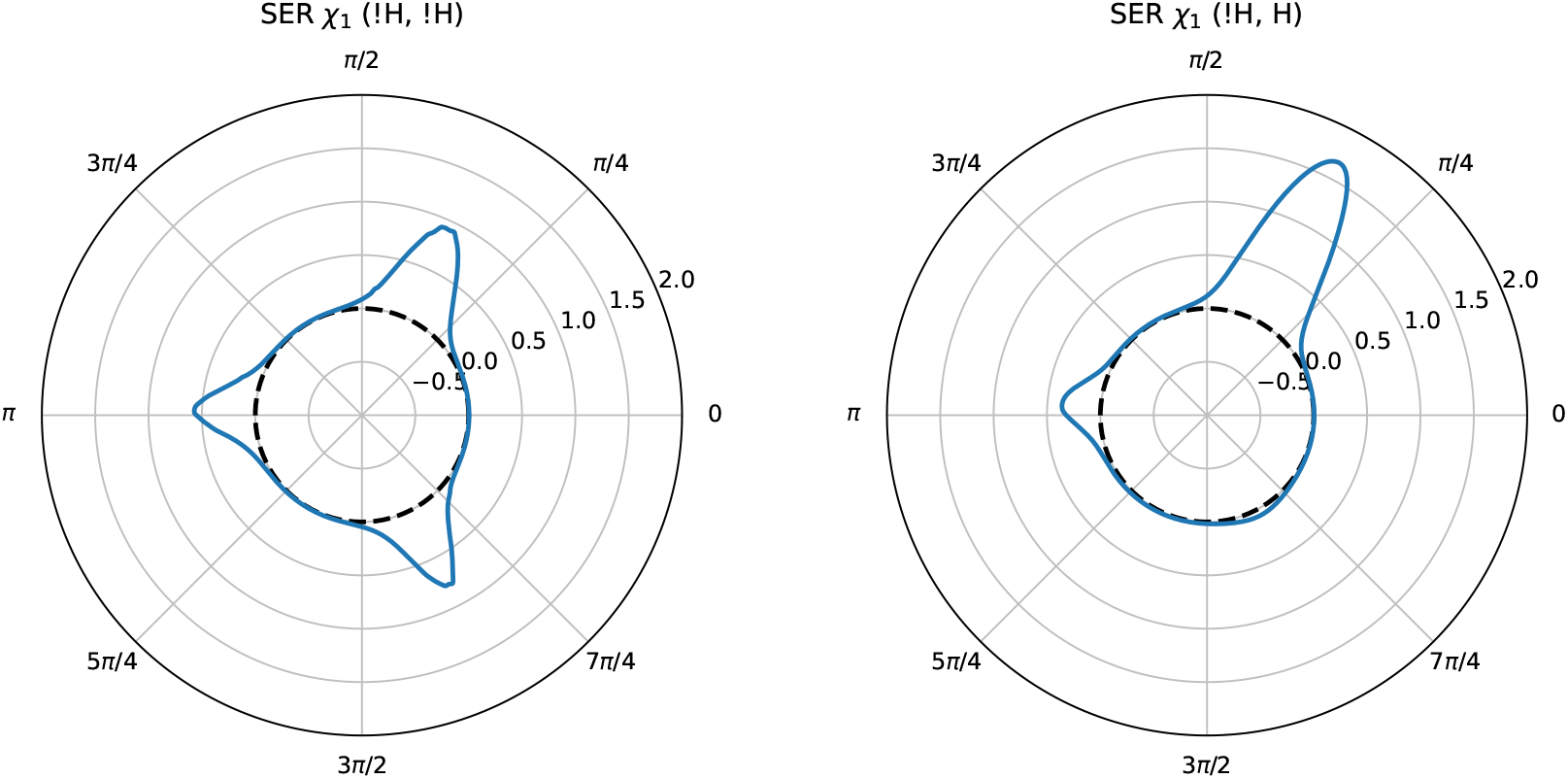
Amino-acid pairs in a specified secondary structure environment: conditioned densities for pairs capture side-chain rotamer preferences in an *α*-helix Ncap environment. Projections of the full torsion-angle probability density for amino-acid pair SER-ALA onto the SER side-chain torsion angle *χ*_1_, conditioned by two different SSE pair labels: *Left:* (! H, ! H), a non-helical environment for both amino acids, and *Right:* (! H, H) consistent with an *α*-helix Ncap environment for SER.

#### Helix transitions and capping

The pertinence of amino-acid pair preferences in the full torsion-angle space for SSE transitions in protein structures can be seen in helix-capping motifs. Helix capping allows stabilization of the ends of *α*-helices by compensating for hydrogen bonds that cannot form at helix termini. For instance, in an infinite regular *α*-helix, the backbone NH group of every residue *i* forms a hydrogen bond with the carbonyl oxygen of residue *I ™* 4. However, in a finite helix, the first four amino acid residues lack these partners. In the N-terminal capping box for a helix starting at residue *i*, the non-helical Ncap residue (*I ™* 1) forms a hydrogen bond with the backbone NH of helical residue *i* + 2. Reciprocally, the side chain of residue *i* + 2 accepts a hydrogen bond from the backbone NH of the Ncap residue, forming a hydrogen-bonded cycle [26]. These interactions partially replace the missing backbone hydrogen bonds and thereby stabilize the helix terminus. Common Ncap residues are Thr, Ser, and Gln, while Glu or Gln are frequent at *i* + 2. We thus note that formation of the capping box thus involves constraints on side-chain *χ* torsion angles.

As an example of the horizons opened by SSE-label conditioned probability densities for amino acid pairs, which are expressed in the full torsion angle space (and which thus include the *χ* side-chain angles), we compare the side-chain *χ*_1_ distribution of the first amino acid in the pair SER-ALA on the two different SSE environments ^2^ { ! Helix - ! Helix } and { ! Helix - Helix}, that is, a non-helical SER followed by a non-helical ALA, and a non-helical SER followed by a helical ALA (this first helical residue is generally referred to as N1). To this end, we specialize the SER-ALA mixture model on both SSE environments, following Eq.2, yielding the *χ*_1_ distribution (Fig. 4). In the non-helical environment { ! Helix - ! Helix}, the SER *χ*_1_ angle distribution indicates the presence of three rotamer modes *g*+, *t*, and *g™* with similar probabilities. However, when preceding a helical environment { ! Helix - Helix}, this distribution is strikingly different: *χ*_1_ rotamer *g*+ is nearly exclusively selected. This reflects the enrichment of Ncap SER residues in the amino-acid pairs capturing the non-Helix to Helix transition: the SER *g*+ rotamer allows the hydrogen-bonding interaction [26] between its side-chain hydroxyl and the backbone amide NH of helical residue N3.

The result just shown is remarkable as well in that the amino-acid pair specification { ! Helix - Helix} that we used to define this SSE-conditioned probability density does not have anything to say about helix position N3 *directly* — indeed N3 is two amino acids further along the helix than the second amino-acid residue of the pair, specified here as the first residue of a helix. But the presence of helical N3 is implicit in DSSP’s SSE definition H used to condition the amino-acid pair densities: only helices containing more than four [8] consecutive helical residues may receive the label H.

## 4 Outlook

The question of understanding the precise relationship between polypeptide chain configurations and protein structure pervades early research in protein science, as reflected in the fundamental work of numerous researchers including Linderstrøm-Lang and Pauling, to cite just two. Since Ramachandran, the interpretation of protein crystal structure data has been facilitated by lower-resolution descriptions in terms of the canonical *φ* and *ψ* backbone torsion angles, which were increasingly associated with hydrogen-bonding analyses, in service of defining the common secondary structures that have furnished a shared vocabulary for discussing the variety of protein structures.

Nevertheless, a quantitative view of the connection between Ramachandran analyses and the resulting secondary structure assignments was lacking. Our definition of SSE-label conditioned torsion-angle densities, founded on the torsion-angle distributions in the full (soft) torsion-angle spaces of the proteins in the current PDB [18], fills this gap. While previous work defining Ramachandran regions and sterically admissible regions had been conducted using a combination of model systems, modeling and data analysis, our systematic processing yields statistically grounded definitions of regions via these conditional densities, which are notably expressed in terms of continuous probability densities as opposed to point clouds.

The new SSE-label conditioned densities allow, first, visual assessment of the correspondence between Ramachandran regions and credible regions of the density for each SSE label, and permit defining an entropy of the SSE labels in Ramachandran space in order to address uncertainty in the assignments. Second, they permit a quantitative assessment of the determinism of SSE labels in terms of backbone torsion angles alone (standard Ramachandran) versus backbone + side chain torsion angles. Finally, the new pair densities – conditioned on the SSE labels for pairs of amino acids as opposed to those for single amino acids – offer new insights. The example we provided of globally analyzing SSE label pairs in SSE transitions in the PDB is just one way of using this approach; the pair densities also revealed their relevance for quantifying more targeted analyses, such as helix capping. Overall, we believe these SSE-conditioned densities offer a compelling framework for quantifying the relationship between Ramachandran diagrams and SSEs in both the single amino acid and amino-acid pair contexts — retaining the simplicity of the classical Ramachandran diagram while encompassing influences arising from correlations in the full torsion angle space.

In terms of applications, we anticipate that our conditioned densities will prove instrumental in multiple tasks. The first is the generation of tuples of torsion angles conditioned on SSE labels. Tuples of torsion angles pulled from the latent space are at the core of the structure module of AlphaFold2. Our conditioned densities, obtained from SOTA statistical models for joint distributions, are certainly a tool of choice to perform such generations. The second impact is in the definition of edge potentials with applications to Ramachandran-aware inverse-kinematics sampling of protein loops and fragments [27], and to free energy estimates based on message passing algorithms [28]. In both cases, our models provide an explicit control on SSE, that remains difficult to enforce explicitly in current AI-based generative models. The third is the reconstruction of protein structures in the absence of homologous templates, a process that can benefit substantially from accurate secondary structure element (SSE) information. However, SSE annotations are often fragmented, with {S, T, C} labels interspersed among {G, H, I} or {B, E} labels. Our novel insights into the coherence between Ramachandran regions and SSE labels provide a unique opportunity to identify and regularize such fragmented regions in terms of SSE labels. Yet another benefit is the design of Goodness-of-fit (GoF) statistical tests, to identify biases inherent to generative engines. Biases in SSE elements have been observed recently by comparing the SSE labels delivered by `DSSP4` on the PDB versus `AlphaFold-DB`. Our conditioned densities provide a direct route to define GoF test, using for example the generic Maximum Mean Discrepancy framework. In turn, such tests should contribute to controlling the biases of generative engines.

## Supporting information

main text

## Funding

This work has been supported by the French government, through the grants: 3IA Côte d’Azur Investments (ANR-23-IACL-0001) (F. Cazals); ANR project Innuendo (ANR-23-CE45-0019) (F. Cazals); ANR-24-RRII-0002 operated by the Inria Quadrant Programme (F. Cazals); Initiative d’Excellence, ANR-11-LABX-0011-01 (C.H. Robert).

## Footnotes

1 https://sbl.inria.fr/data/higher-ramachandran/

2 The notation !H refers to not Helix

## Notes

### Competing Interest Statement

The authors have declared no competing interest.

