## Supplementary material for "Beyond Ramachandran distributions: quantifying secondary structure effects on amino acids and their pairs in the full torsional space": main text

### S1 SSE: refresher

Refresher for Fig. 1.

**DSSP4 classification for Secondary Structure Elements (SSE).** The classification of SSE in DSSP4 defines 9 labels as follows [9]:

- Based on H-bonding patterns:
  - (Labels G/$3_{10}$-helix, H/$\alpha$-helix, I/$\pi$-helix) Three types of helices, named after the number of residues skipped to form the H-bond, that is $(i, i+3)/(i, i+4)/(i, i+5)$.
  - (Label P/$\pi$-helix) Also named PPII-helix– see below.
  - (Label T/Turn) Hydrogen binding pattern if 3/4/5 helix not already assigned.
  - (Labels B/$\beta$-bridge, E/extended $\beta$ strand) Two types of $\beta$ structures which can be parallel or anti-parallel. $\beta$-bridges are isolated, while repeated $\beta$-bridges form extended $\beta$-strands.
- Without any specific bonding pattern:
  - (S/Bend) Characterized by $\angle C_{\alpha;i-2}C_{\alpha;i}, C_{\alpha;i}C_{\alpha;i+2} \geq 70°$
  - (C) No other label assigned: termed Coil in the sequel.

**H bonds.** We may also recall classical hydrogen bonding patterns. For say a label H / 4-helix, the hydrogen bonding pattern is is CO(i)-NH(i+4), with CO(i) acceptor and NH(i+4) donor. For label T, the hydrogen-bonded turn generally corresponds a H bond $CO(i)$ and $NH(i+n)$ not already assigned to a helix [24, 13], with $n = 2$ for a $\gamma$-turn and $n = 3$ for a $\beta$-turn [29] – the cases $n = 3, 4, 5$ corresponding to helices.

**Ramachandran plot nomenclature (from [13]).** Empirically, one notes the presence of relatively homogeneous Ramachandran regions:

1. region $\alpha$ for helices (mostly DSSP4 labels: GHI).
2. region $\beta$ for strands (mostly DSSP4 labels: BE).
3. region $\kappa$ pour PPII helices (mostly DSSP4 label: P).
4. region $\gamma, \gamma'$: the H-bond $CO(i) \rightarrow NH(i+2)$. Mostly the DSSP4 label $T$.

and more complex regions:

1. region $\delta$: the slanted region above the $\alpha$ region.
2. specific region $\varepsilon$: largely populated by GLY (no $C_\beta$ carbon, see below).
3. specific region $\zeta$: specific to residues preceding PRO;
4. specific region $\varepsilon$: vastly populated by GLY due to the absence of $C_\beta$ atom.

This heterogeneity indeed attests to the difficulties of interpreting secondary structure in the Ramachandran space alone.

### S2 SSE conditioned densities

#### S2.1 Theory

We prove proposition 1.

**Samples and associated indices.** To fit the densities, we consider pairs $(x, z)$ with $x \in \mathbb{T}^d$ and $z \in 1 \dots K$ the latent variable corresponding to the index of the mixture component which generated $x$.

We associate to each mixture component $C_k$ a latent variable $D_k$ taking values the dictionary $\mathcal{L}$ of SSE labels. Note that in practice, this variable is associated to the function $\texttt{label}(x) \in \mathcal{L}$ returning the label of a point/an observation $x$.

**Responsibility of a mixture for a sample.** The *responsibility* $r_k(x)$ of each component in a mixture for a sample $x$ is the following posterior probability [30]:

$$r_k(x) = \mathbb{P}\left[z = k \mid x\right] = \frac{\mathbb{P}\left[x \mid z = k\right]\mathbb{P}\left[z = k\right]}{\mathbb{P}\left[x\right]} = \frac{w_k f_k(x)}{\sum_{j=1}^{K} w_j f_j(x)}. \tag{3}$$

Note that $\sum_k r_k(x) = 1, \forall x$.

**Responsibility of a label for a component.** Consider the responsibilities of the labels for the k-th component, namely the vector $(m_{k1}, \dots, m_{kS})^{\mathsf{T}}$ – with $\sum_{j \in \mathcal{L}} m_{kj} = 1, \forall k$.

Using Bayes' formula yields

$$m_{kj} = \mathbb{P}\left[D_k = j \mid z = k\right] \tag{4}$$

$$= \frac{1}{w_k} \int_{\mathbb{T}^d} \mathbb{P}\left[z = k \mid x\right] \mathbf{1}_{\texttt{label}(x)=j}\, p(x) dx \tag{5}$$

$$= \frac{1}{w_k} \int_{\mathbb{T}^d} r_k(x) \mathbf{1}_{\texttt{label}(x)=j}\, p(x) dx. \tag{6}$$

Let $X_{|j}^{\text{Data}}$ be the subset of $X$ with label $j$. The previous yields the estimator

$$\hat{m}_{kj} = \frac{\sum_{i \in X_{|j}^{\text{Data}}} r_k(x_i)}{\sum_i r_k(x_i)}. \tag{7}$$

**Probability of a label.** The unconditional probability that the mixture generates a label $j \in \mathcal{L}$ is

$$\mathbb{P}\left[\texttt{label} = j\right] = \sum_k \mathbb{P}\left[D_k = j | z = k\right] \mathbb{P}\left[z = k\right] = \sum_k w_k m_{kj}. \tag{8}$$

**Proof of proposition 1.** Using the previous ingredients, we prove the following expression from proposition 1:

$$p(x \mid \texttt{label}(x) = j) = \sum_k \frac{m_{kj} w_k f_k(x)}{\sum_k m_{kj} w_k}.$$

*Proof.* Let us write

$$p(x, \texttt{label} = j) = \sum_k p(x, \texttt{label} = j, z = k). \tag{9}$$

Assuming that the label reweights but does not change the functional expression of the components:

$$p(x, \texttt{label} = j, z = k) = p(x | \texttt{label} = j, z = k) p(\texttt{label} = j | z = k) p(z = k) \tag{10}$$

$$= f_k(x) m_{kj} w_k. \tag{11}$$

But using Eq. 8 yields:

$$p(x | \texttt{label} = j) = \frac{p(x, \texttt{label} = j)}{\mathbb{P}\left[\texttt{label} = j\right]} \tag{12}$$

$$= \frac{\sum_k f_k(x) m_{kj} w_k}{\sum_k m_{kj} w_k}. \tag{13}$$

□

The estimator for the previous is obtained using the expression of $\hat{m}_{kj}$ in Eq. (7).

**SSE label uncertainty in Ramachandran space via Shannon's entropy.** With $\mathcal{L}$ the set of SSE labels, given a sample of size $n$, the entropy of SSE labels in a bin of the Ramachandran map reads as

$$H(X, \mathcal{L}) = -\sum_{j \in \mathcal{L}} p(j) \ln p(j). \tag{14}$$

Which we estimate as follows

$$\tilde{H}(X, \mathcal{L}) = -\sum_{j \in \mathcal{L}} \frac{\sum_{i=1}^{n} \mathbf{1}\{z_i = j\}}{n} \ln \left( \frac{\sum_{i=1}^{n} \mathbf{1}\{z_i = j\}}{n} \right).$$

Note that the maximum entropy corresponds to a uniform distribution on labels and is equal to $\ln S$.

**Remark 1** *To render the entropy map, we proceed as follows: the pixel color is chosen using a viridis color map. The alpha channel (transparency) is used to encode the abundance of samples is that pixel: its value is 1 for $n > 100$, and is linearly interpolated between 0 and 1 otherwise. Thus, no data in a pixel yields a transparent pixel.*

**Jensen-Shannon divergence between conditional distributions on SSE.** Recall that the Jensen-Shannon divergence is a symmetric measure of dissimilarity between two probability distributions derived from the Kullback-Leibler divergence [23]. Let $m(x) = (p(x) + q(x))/2$. The Jensen-Shannon divergence between two distributions $p(x)$ and $q(x)$ reads as

$$D_{\mathrm{JS}}(p \| q) = \frac{1}{2} D_{\mathrm{KL}}(p \| m) + \frac{1}{2} D_{\mathrm{KL}}(q \| m), \tag{15}$$

with the Kullback-Leibler divergence is $D_{\mathrm{KL}}$ defined by

$$D_{\mathrm{KL}}(p \| q) = \int p(x) \ln \frac{p(x)}{q(x)} dx. \tag{16}$$

In practice, we estimate the expectation of $D_{\mathrm{KL}}$ using Monte Carlo sampling. Given samples $X \sim p$ with $|X| = N_p$ and $Y \sim q$ with $|Y| = N_q$, the estimator is

$$\widehat{D}_{\mathrm{JS}}(p \| q) = \frac{1}{2N_p} \sum_{i=1}^{N_p} \ln \frac{p(x_i)}{m(x_i)} + \frac{1}{2N_q} \sum_{i=1}^{N_q} \ln \frac{q(y_i)}{m(y_i)} \tag{17}$$

For our application, we assume that the PDB50 samples used to condition the mixture models are representative of the conditioned models. As such, to compute $\widehat{D}_{\mathrm{JS}}$ between conditional mixtures on SSE labels $i$ and $j$, we take $X = X_{|i}^{\mathrm{Data}}$ and $Y = X_{|j}^{\mathrm{Data}}$.

### S3 Artwork

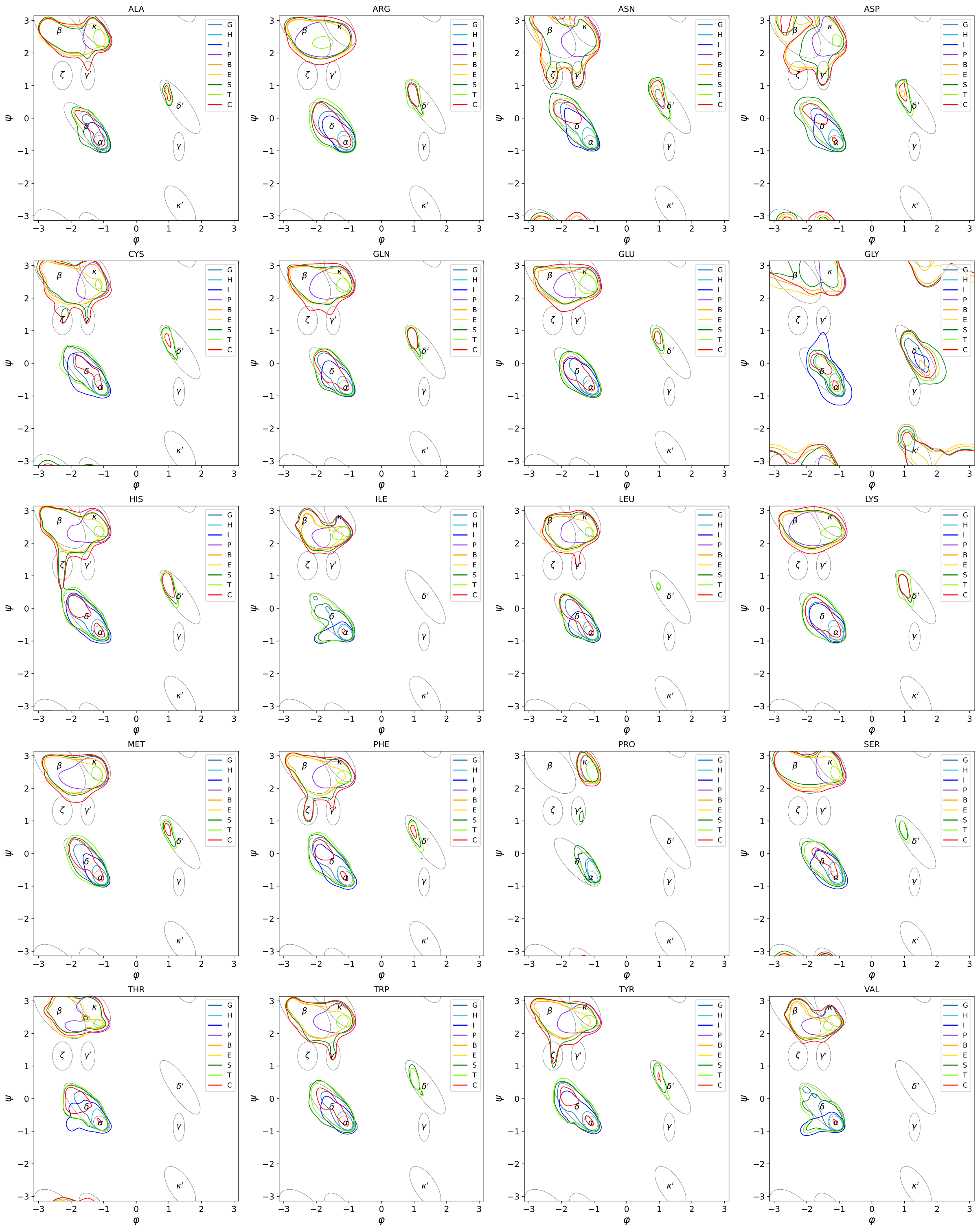

Figure S1: **Densities of specialized submodels superposed with Ramachandran regions.** Credible region contour at 75% level.

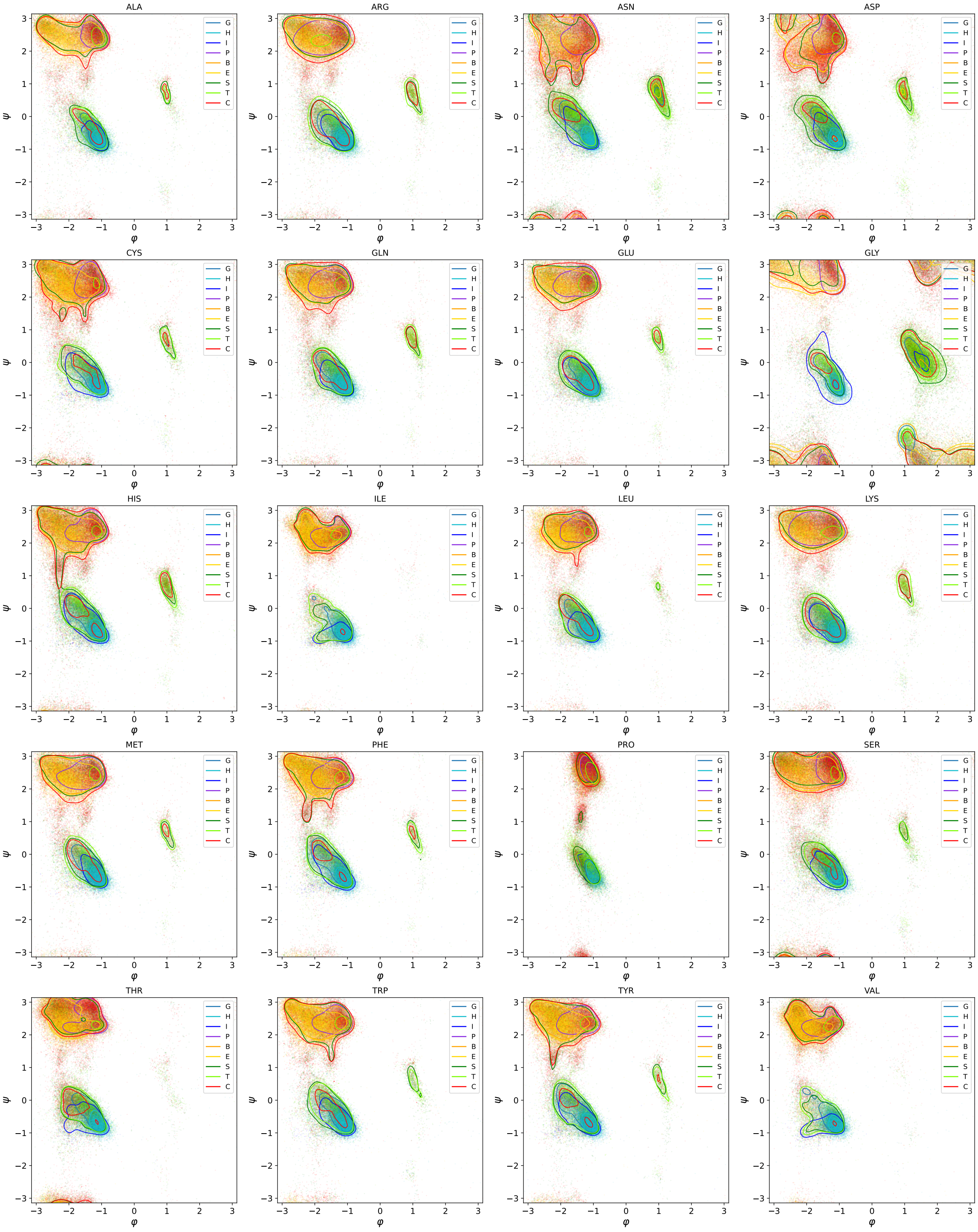

Figure S2: **Samples of 100,000 datapoints for each amino acid superposed with credible regions of specialized submodels densities at 75% level.**

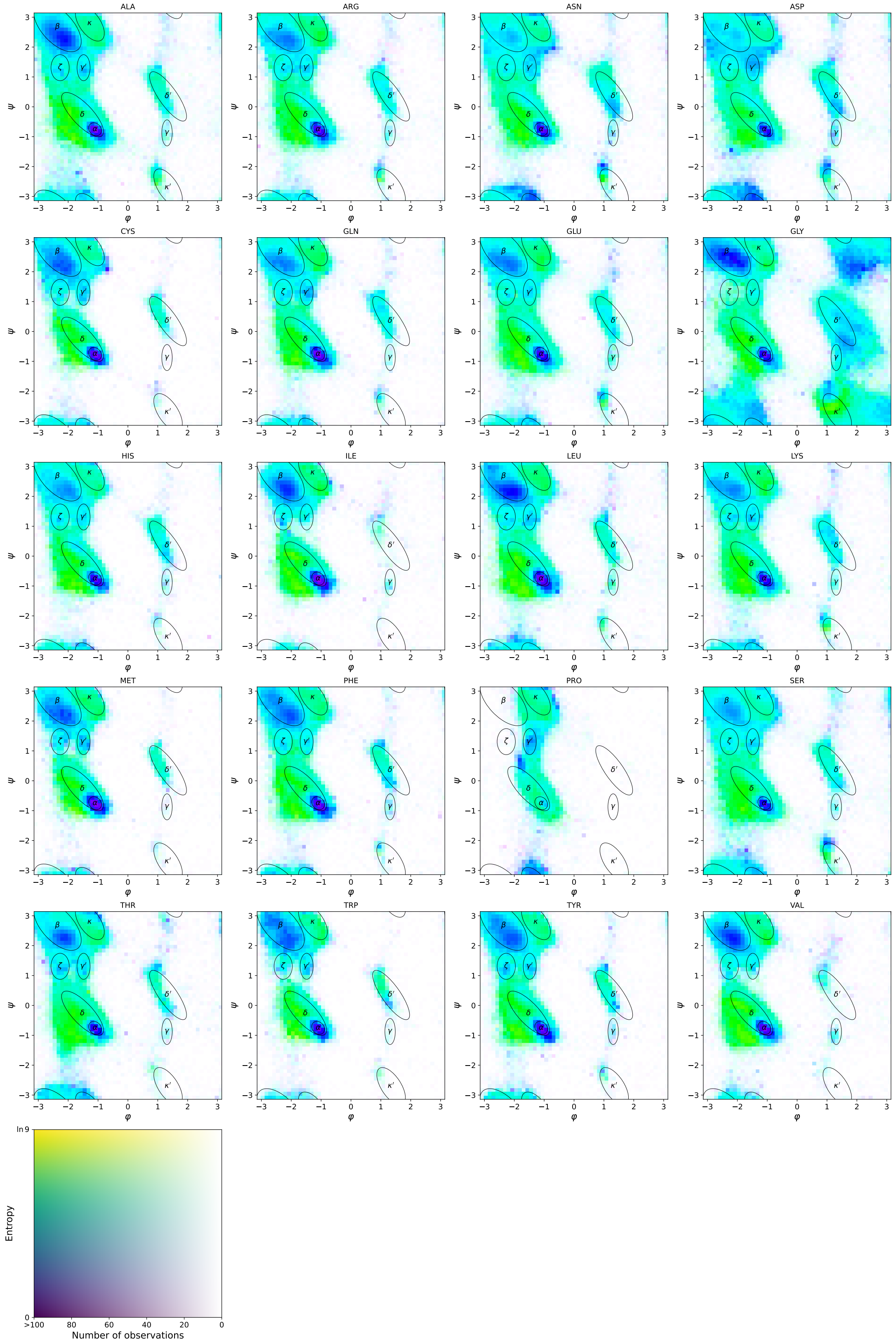

Figure S3: **Entropy in DSSP4 classification based on Ramachandran position.** Entropy is estimated for each bin in a 25x25 grid using Eq. 14. Entropy values range from minimum entropy at 0, to max entropy at ln(9) – uniform distribution over 9 DSSP4 labels. Transparency is proportional to the amount of observations at given cell – ranging from minimum at 0, to maximum at 100 observations and above

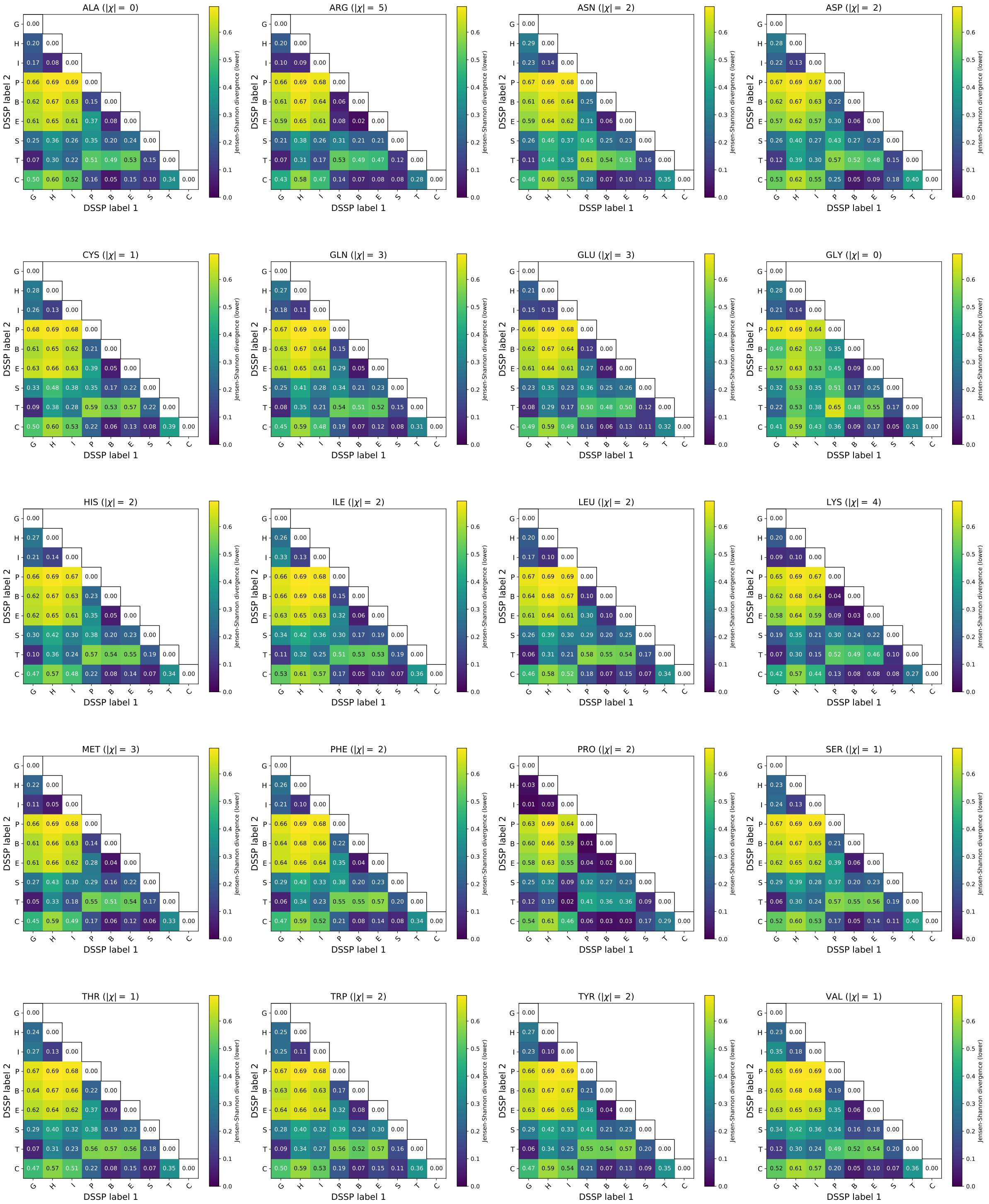

Figure S4: **Jensen-Shannon divergence estimates between conditioned models: per amino acid using all torsion angles.** Colormap scaled using the absolute range $[0, \ln(2) \approx 0.69]$;

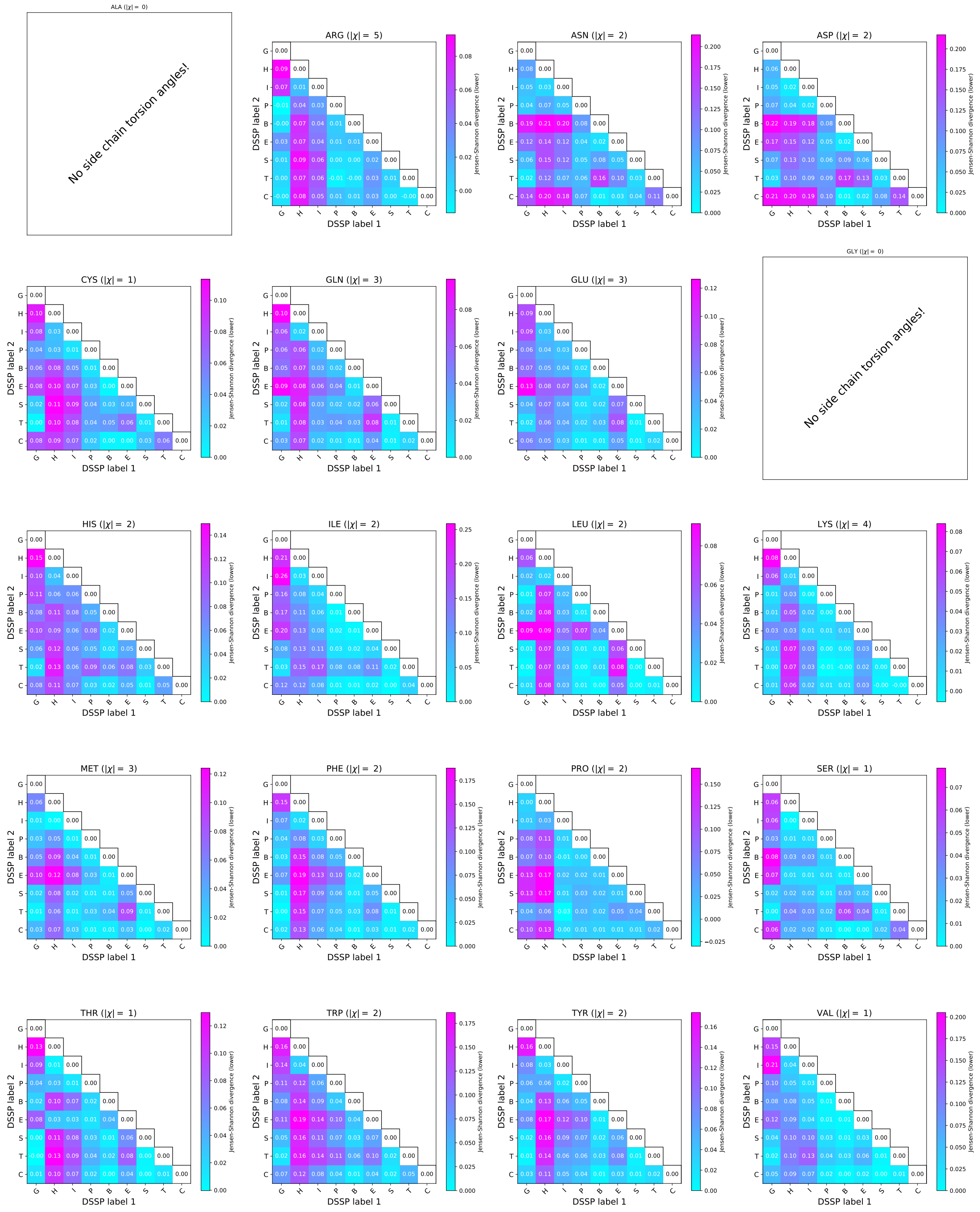

Figure S5: **Jensen-Shannon divergence estimates between conditioned models: per amino acid using Side-chain torsion angles only.** Colormap scaled using the range 0 - max value observed.

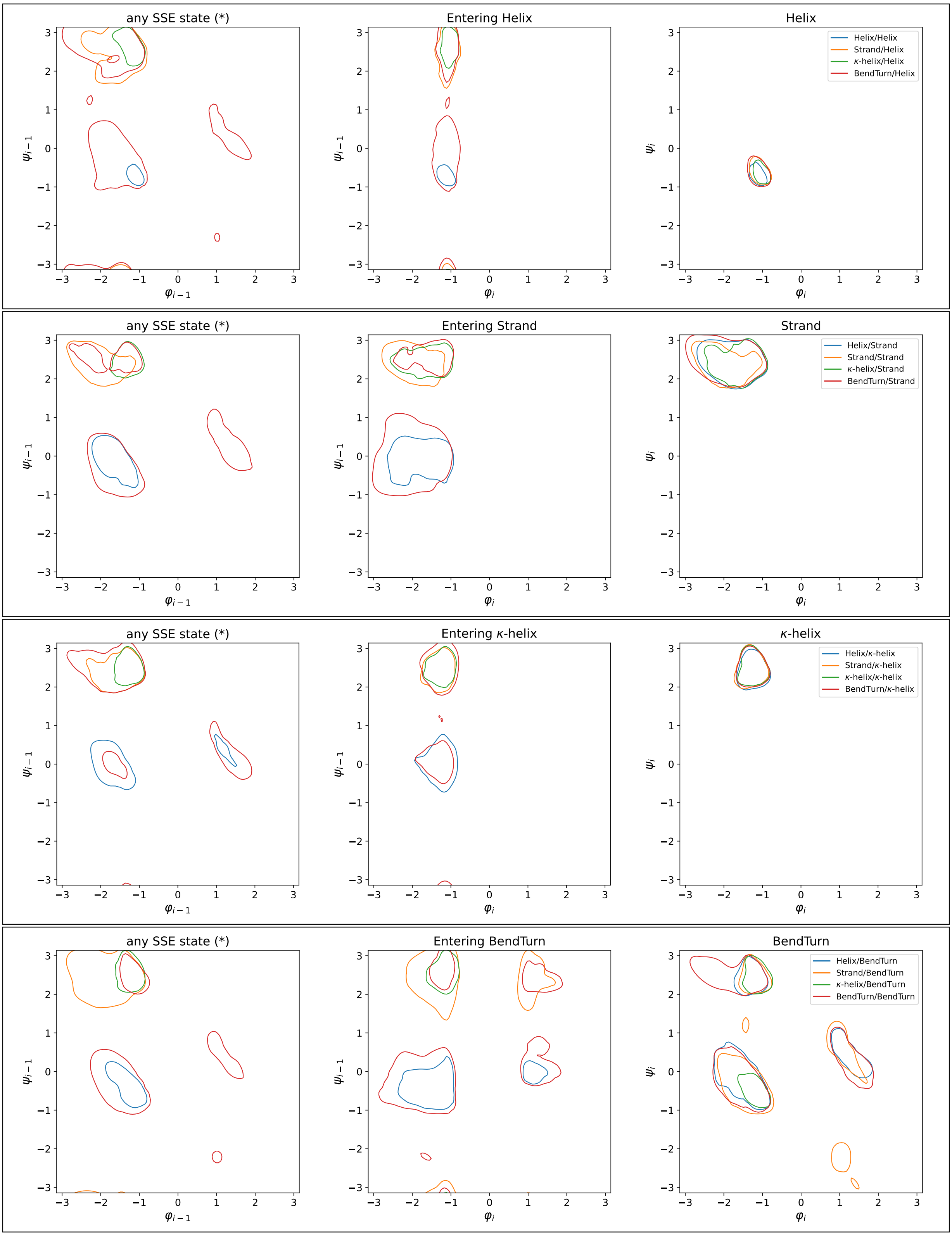

Figure S6: **Entering a SSE: densities of specialized submodels in amino acid pairs.** All amino acids considered. Helix combines the {'H','G','I'} DSSP labels, Strand combines {'B','E'}, $\kappa$Helix is {'P'}, and BendTurn combines {'S','T'}. Credible region contour at 75% level.

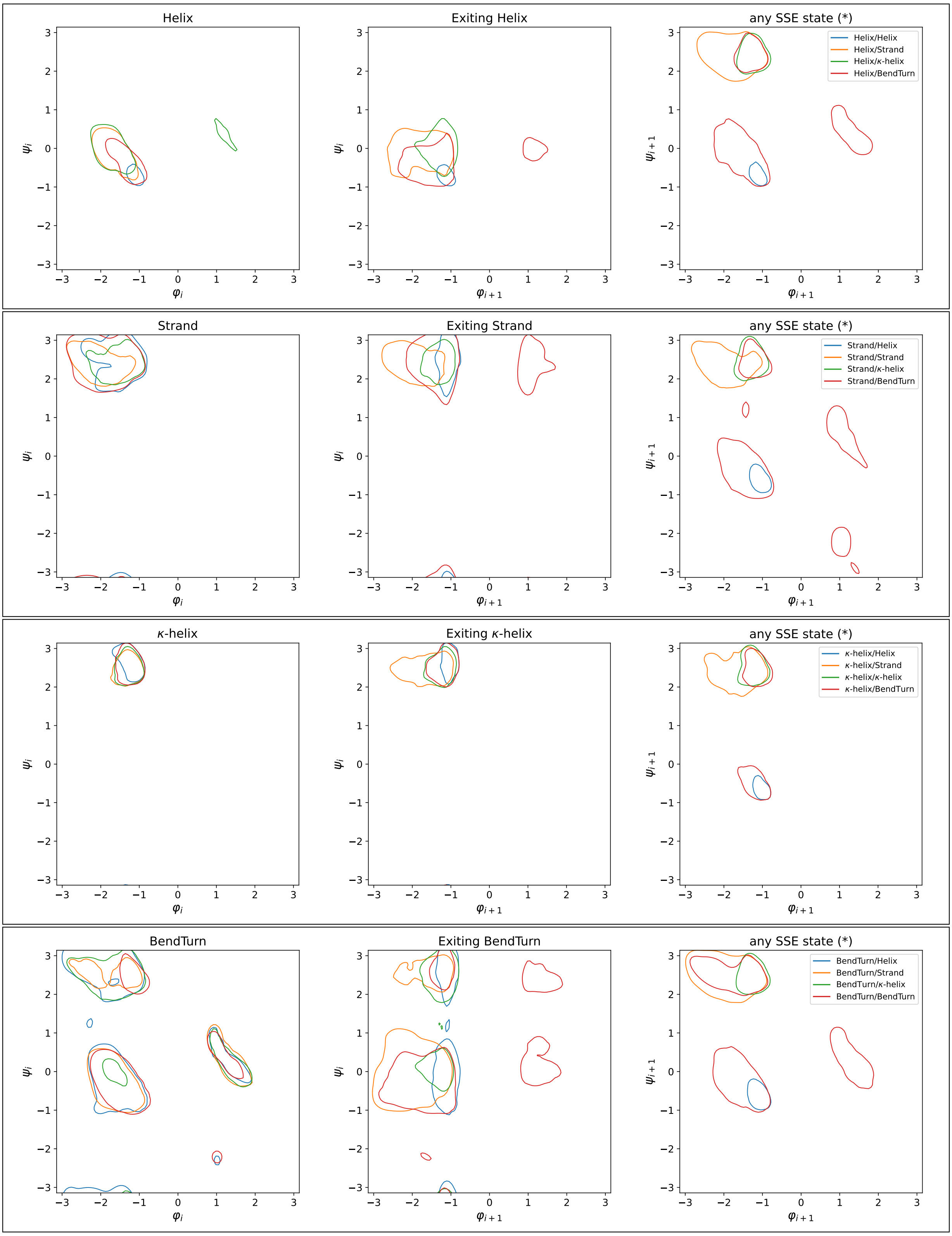

Figure S7: **Exiting a SSE: densities of specialized submodels in amino acid pairs.** All amino acids considered. C Helix combines the {'H','G','I'} DSSP labels, Strand combines {'B','E'}, $\kappa$Helix is {'P'}, and BendTurn combines {'S','T'}. Credible region contour at 75% level.

### S4 Specialized submodel densities

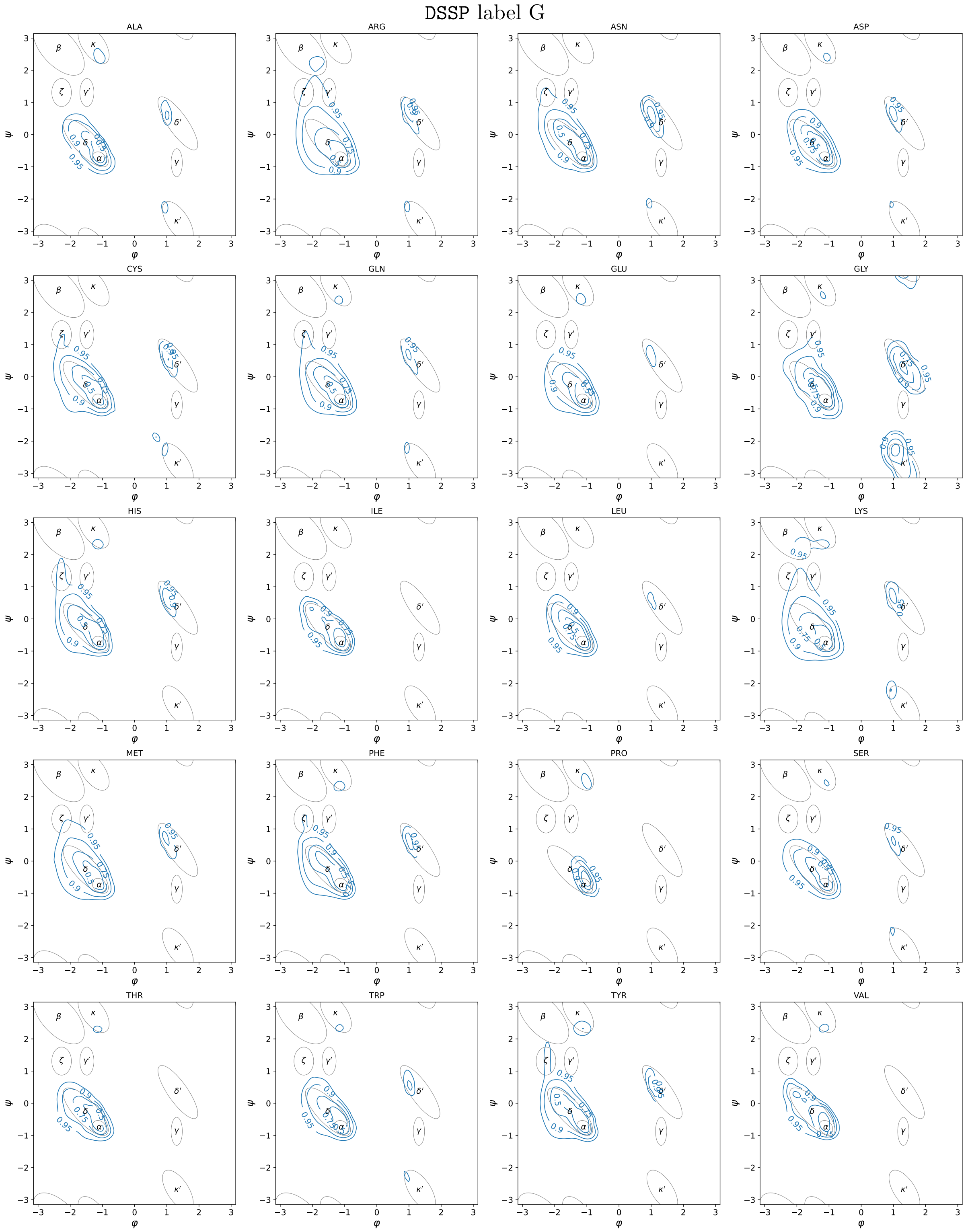

Figure S8: **Density of the submodel specialized on 'G' DSSP label.** Credible region contours at thresholds {95%, 90%, 75%, 50%}

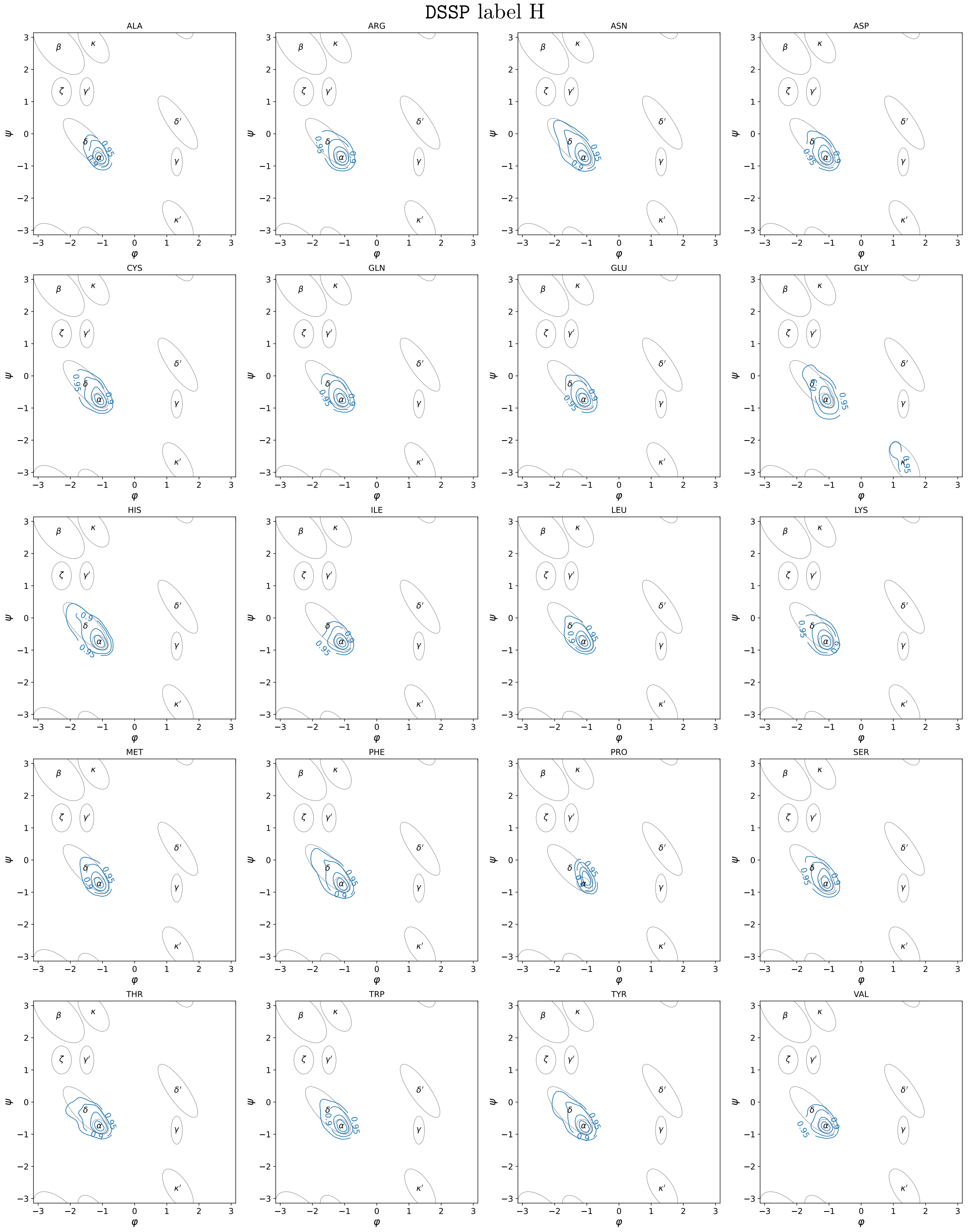

Figure S9: **Density of the submodel specialized on 'H' DSSP label.** Credible region contours at thresholds {95%, 90%, 75%, 50%}

DSSP label I

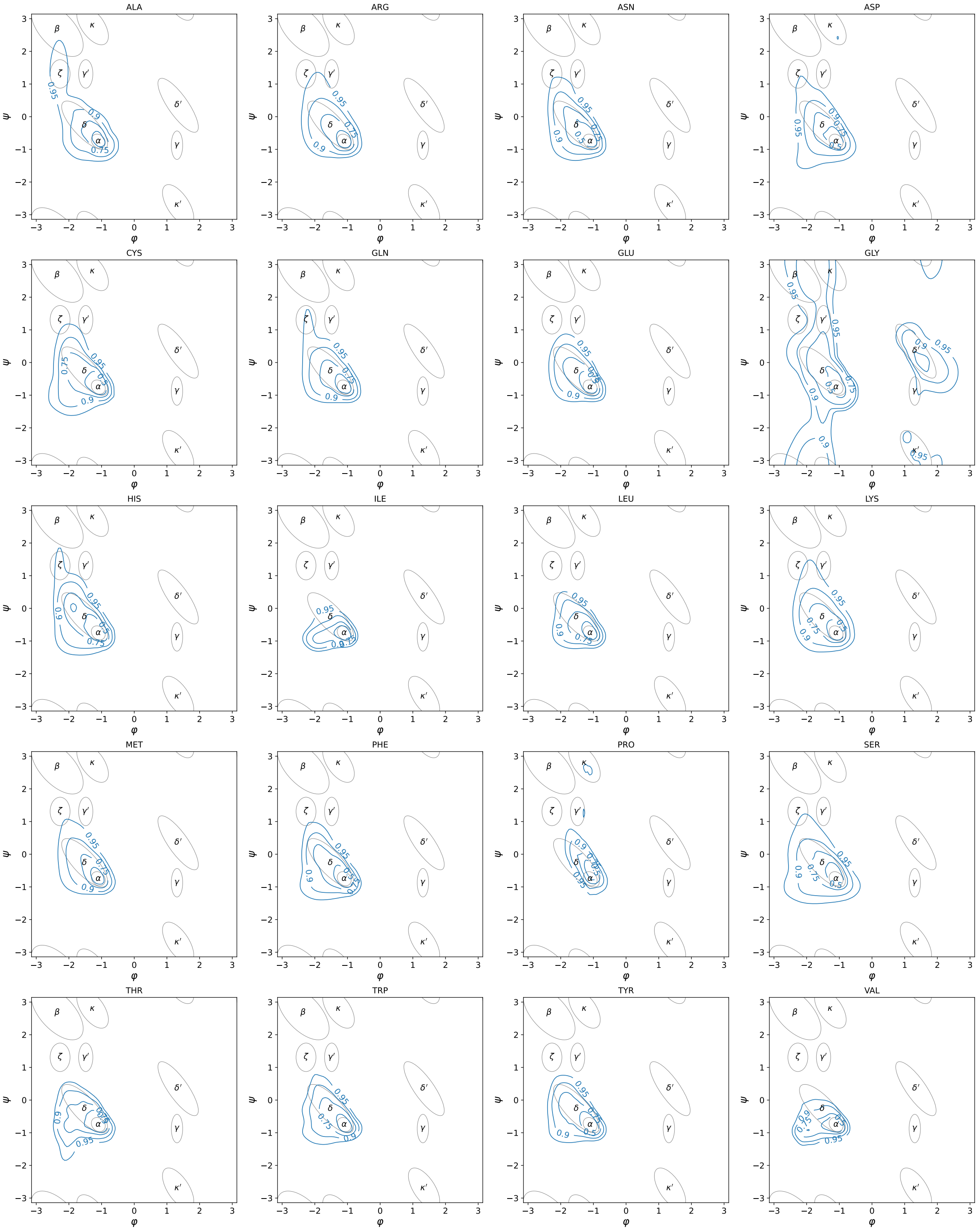

Figure S10: **Density of the submodel specialized on 'I' DSSP label.** Credible region contours at thresholds {95%, 90%, 75%, 50%}

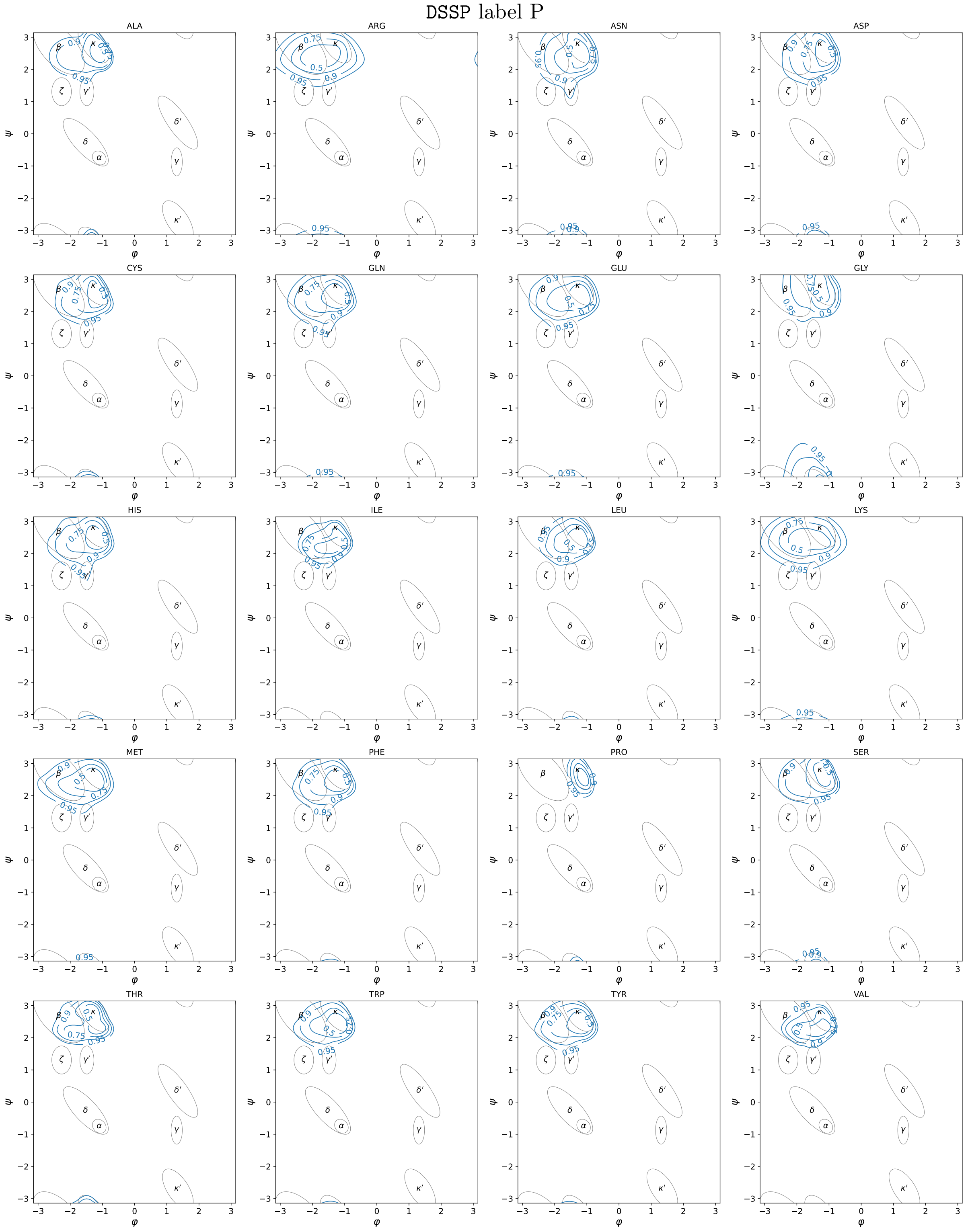

Figure S11: **Density of the submodel specialized on 'P' DSSP label.** Credible region contours at thresholds {95%, 90%, 75%, 50%}

#### DSSP label B

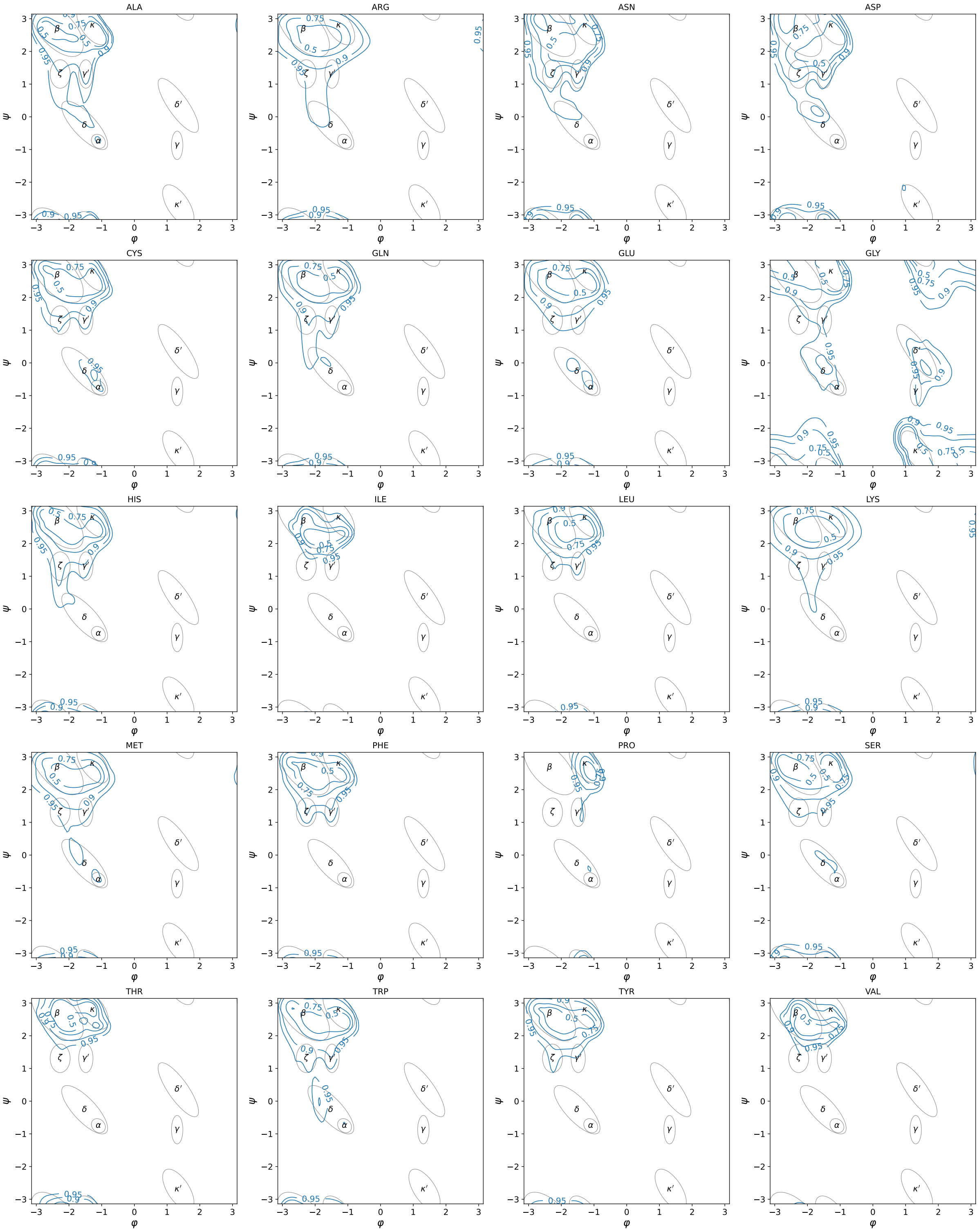

Figure S12: **Density of the submodel specialized on 'B' DSSP label.** Credible region contours at thresholds {95%, 90%, 75%, 50%}

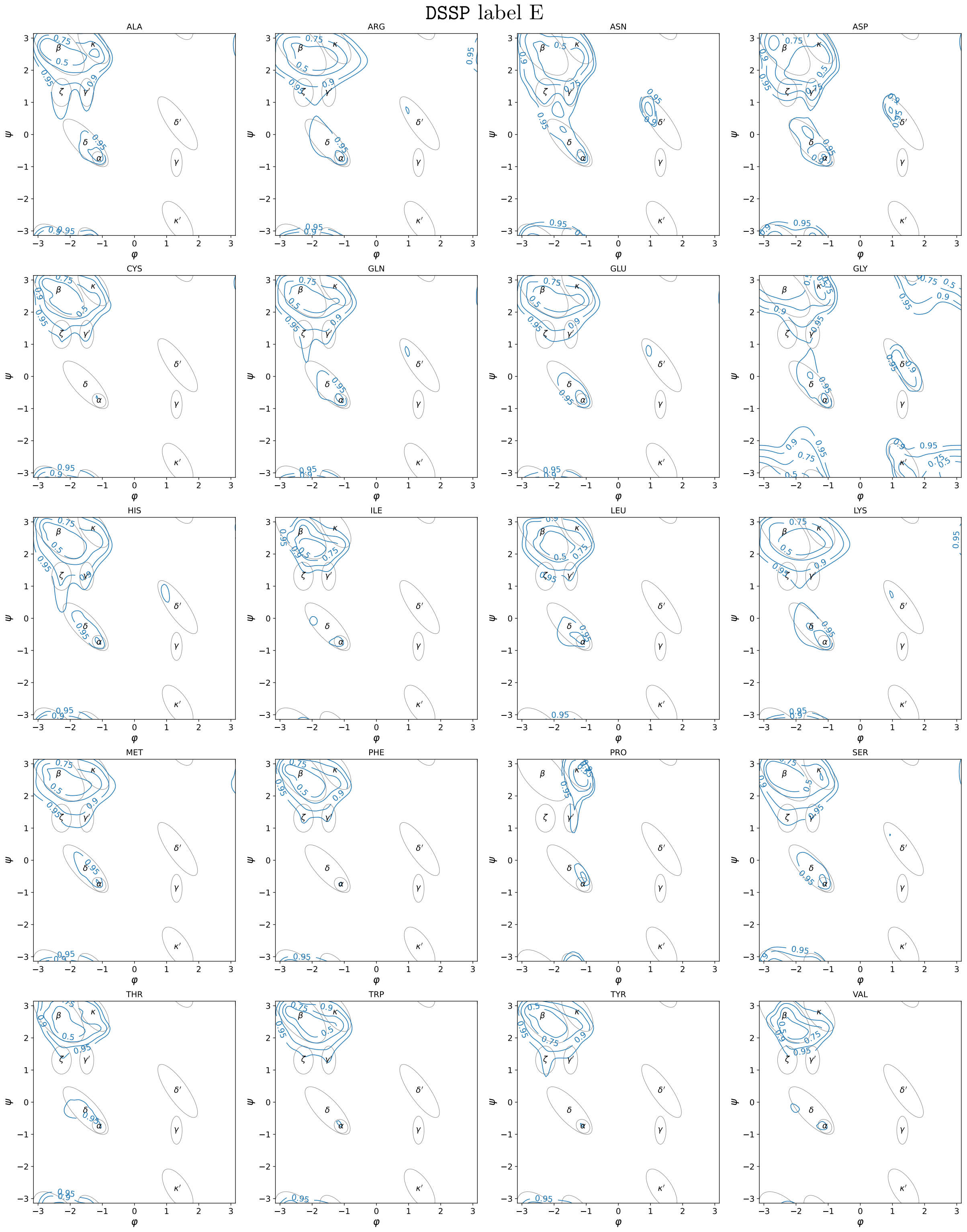

Figure S13: **Density of the submodel specialized on 'E' DSSP label.** Credible region contours at thresholds {95%, 90%, 75%, 50%}

DSSP label S

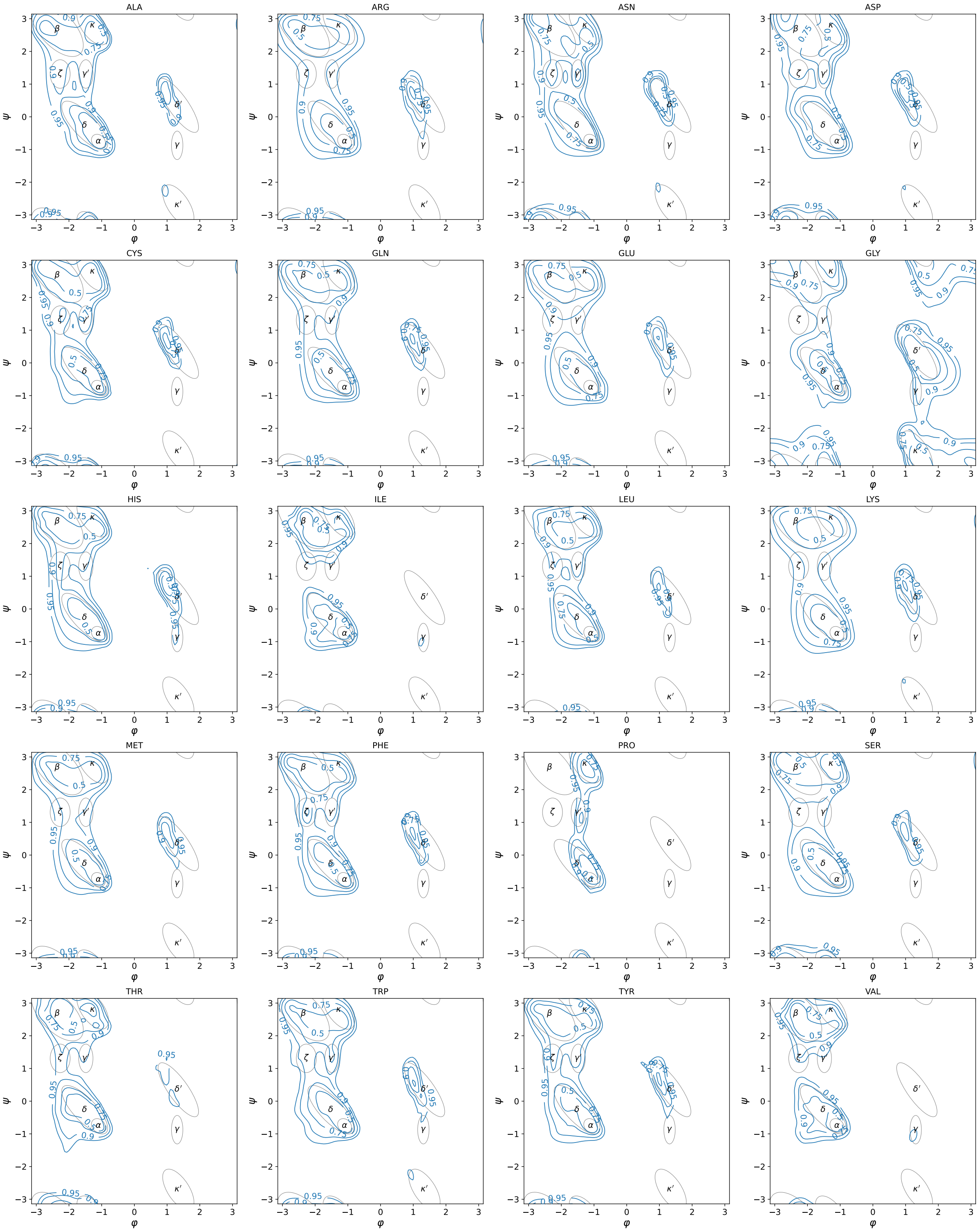

Figure S14: **Density of the submodel specialized on 'S' DSSP label.** Credible region contours at thresholds {95%, 90%, 75%, 50%}

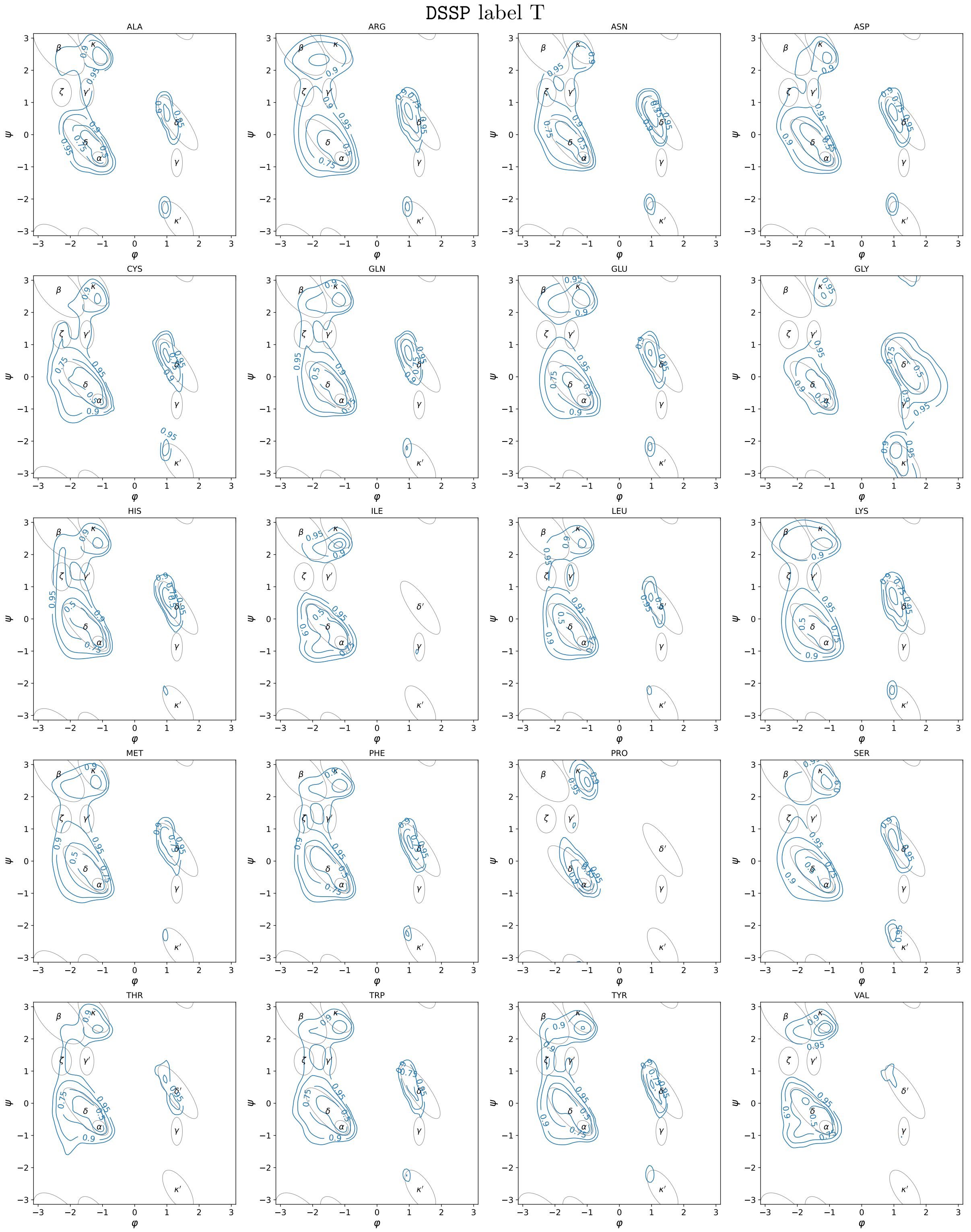

Figure S15: **Density of the submodel specialized on 'T' DSSP label.** Credible region contours at thresholds {95%, 90%, 75%, 50%}

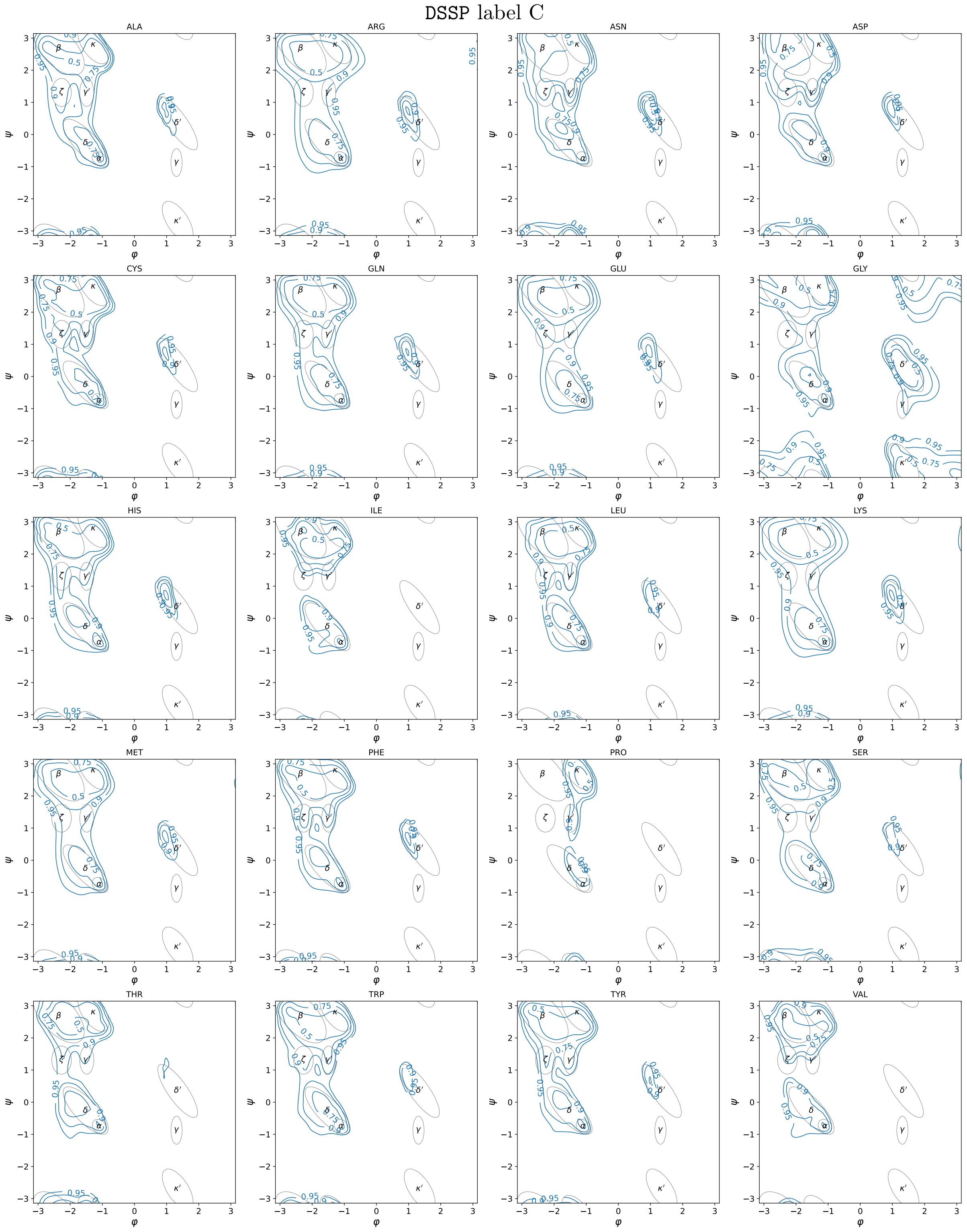

Figure S16: **Density of the submodel specialized on 'C' DSSP label.** Credible region contours at thresholds {95%, 90%, 75%, 50%}

**4 Outlook** **8**

**S1 SSE: refresher** **14**

**S2 SSE conditioned densities** **14**
S2.1 Theory . . . 14

**S3 Artwork** **16**

**S4 Specialized submodel densities** **24**
